# Reward modulates cortical representations of action

**DOI:** 10.1101/2020.01.15.907006

**Authors:** Tyler J. Adkins, Taraz G. Lee

**Affiliations:** Department of Psychology, University of Michigan, Ann Arbor, MI, 48109

## Abstract

People are capable of rapid on-line improvements in performance when they are offered a reward. The neural mechanism by which this performance enhancement occurs remains unclear. We investigated this phenomenon by offering monetary reward to human participants, contingent on successful performance in a sequence production task. We found that people performed actions more quickly and accurately when they were offered large rewards. Increasing reward magnitude was associated with elevated activity throughout the brain prior to movement. Multivariate patterns of activity in these reward-responsive regions encoded information about the upcoming action. Follow-up analyses provided evidence that action decoding in pre-SMA and other motor planning areas was improved for large reward trials and successful action decoding was associated with improved performance. These results suggest that reward may enhance performance by enhancing neural representations of action used in motor planning.

**Highlights:**

- Reward enhances behavioral performance.
- Reward enhances action decoding in motor planning areas prior to movement.
- Enhanced action decoding coincides with improved behavioral performance.

## Introduction

People perform better when they are highly motivated. In experimental settings, motivation has been shown to enhance motor skills (Wächter et al. 2009; Mosberger et al. 2016), working memory (Krawczyk et al. 2007; Kennerley and Wallis 2009), attention (Engelmann and Pessoa 2007), cognitive control (Etzel et al. 2016), model-based decision-making (Patzelt et al. 2018), and perception (Serences 2008). Motivation is often manipulated using prospective cues that involve offers of future monetary gains or threats of future losses. People are capable of rapid improvements in skill performance when they are presented with large prospective incentive cues. This capacity suggests that people update their expectations of future reward prospectively, leading to more attention and effort being invested in the task, and ultimately increasing the probability of success (Kool and Botvinick 2018).

Motor skill learning is sometimes studied in the context of motor sequencing tasks such as the serial reaction time task (SRTT) or the Discrete Sequence Production (DSP) task (Hikosaka et al, 2002; Diedrichsen and Kornyasheva, 2015; but see Krakauer et al. 2019 for more on the role of sequence learning in skill learning). In these tasks, participants train to quickly and accurately perform specific sequences of finger movements. Early on in skill learning, performance is slow, effortful, and unstructured and appears to rely heavily on cognitive processes such as attention and working memory that can support intentional motor planning. This is evidenced by the susceptibility of novice skills to distraction effects (Nissen and Bullemer 1987), the dependency of skill acquisition on working memory capacity (Anguera et al. 2011; Seidler et al. 2012), and the increased activity in brain regions important for cognitive control such as lateral prefrontal cortex (LPFC) during the performance of novice skills (Grafton et al. 1995; Willingham et al. 2002; Schendan et al. 2003). A recent study using the DSP task showed that the performance of explicitly trained (cued) sequences was enhanced more by reward than the performance of implicitly trained (uncued) sequences (Anderson et al., 2020). Since advanced planning was possible only for the cued sequences, this result suggests that reward enhances performance through cognitive control processes. In the present study, we examine the specific hypothesis that reward enhances representations of action used in motor planning.

Several studies show that motor skills can be decoded from patterns of brain activity using multivariate techniques such as pattern classification and representational similarity analysis. For example, sequential actions can be decoded from brain activity before and during movement (Wiestler and Diedrichsen 2013; Nambu et al. 2015) and representational similarity analysis has recently been used to model the structure of motor-skill representations in a motor planning hierarchy spanning pre-supplementary motor area (pre-SMA) to M1 (Yokoi and Diedrichsen 2019). In the context of a cognitive control task, one recent study showed that motivation enhances temporary goal representations in the prefrontal cortex and that this enhancement mediated the effect of motivation on performance (Etzel et al. 2016). This work suggests that increased motivation may also enhance motor skill representations, particularly if their successful performance depends on cognitive control. We therefore expected that the motivated performance of motor skills would be accompanied by improvements in the decodability of action information in the lateral prefrontal cortex and higher-level motor planning areas. No study to date has examined this possibility.

We administered a discrete sequence production (DSP) task during functional magnetic resonance imaging (fMRI) to 30 healthy human participants. Participants trained for 40 trials on one sequence and 200 trials on another and returned 48 hours later to perform these sequences for prospective rewards ($5, $10, or $30). We found that behavioral performance was enhanced for $30 trials compared to $5 trials. Our data revealed a widespread network of brain areas whose activity scaled linearly with reward magnitude. From this reward-responsive network, we decoded information about upcoming action and performance from patterns of activity preceding movement. This distributed representation included clusters in movement planning areas such as LPFC, pre-supplementary motor area (pre-SMA) and supplementary motor area (SMA). We then examined whether action decoding in specific ROIs was influenced by reward magnitude. We found that action decoding in pre-SMA was enhanced for trials with large reward cues. Furthermore, decoding in SMA was associated with improvements in behavioral performance. Although future work is needed to determine exactly what aspects of action—e.g., sequence, vigor, skill level—are encoded in these brain areas, our results suggest that reward may improve performance by enhancing action coding prior to movement.

## Materials and Methods

### Participants

30 undergraduate students from the University of Michigan participated in this study. All participants gave written informed consent to participate and were compensated at a rate of $15 per hour plus cash bonuses for successful performance. All materials and methods were approved by the University of Michigan Institutional Review Board.

### Behavioral task

Participants learned to perform two sequences of eight keypresses (Fig. 1). Before each trial, participants were instructed by a color cue (1.5 s) to perform one of the two sequences. After a brief delay (2-6 s), participants could perform the instructed sequence using the A-S-D-F keys. During movement, an array of 4 grey rectangles (representing the keys) ‘lit up’ in sequence to remind the participant of which key to press next. If an incorrect key was pressed, the corresponding placeholder box would turn red for one second and the trial was aborted. If the participant did not successfully complete the entire eight-item sequence under the timed deadline (see below), a message saying “Too Slow” was displayed for one second and the trial was aborted. There was no feedback following a successful trial. We expected that our participants would rely less on these on-line visual cues, and more on advanced planning, as learning progressed. In the reward session, sequence cues were presented concurrently with incentive. Stimulus orderings for sequence identity and the duration of cue-to-execution intervals (2-6 s) and inter-trial intervals (2-6s) were optimized to estimate effects of interest using AFNI’s make_random_timing tool by running 5000 iterations of different random orderings/timings and selecting those explained the most variance in simulated GLM analyses.

**Figure 1.**
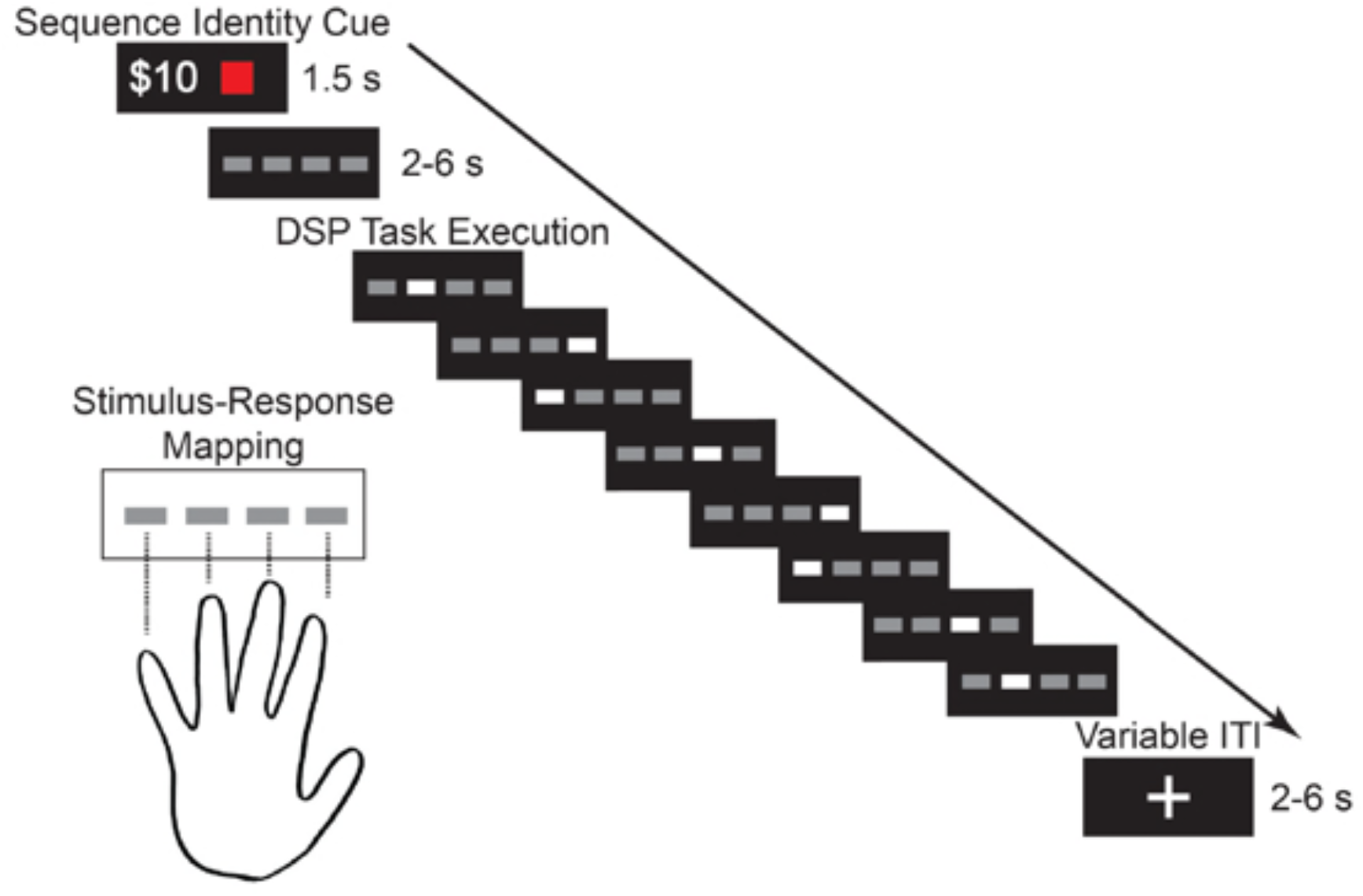
Discrete sequence production task. To study the effects of motivation on skilled action, participants performed two 8-item motor sequences for monetary incentives. The diagram above depicts an example trial from the reward session (training was the same except there were no incentives). Reward and sequence (color) were cued at the start of each trial. Our fMRI analyses focused on hemodynamic responses to this cue. After a brief delay, the sequence was performed. If it was completed under a specific time limit, the trial was successful.

### Experimental Protocol

Participants performed the DSP task described above during fMRI scanning in two separate sessions 48 hours apart. In session one, participants performed eight blocks of 35 trials each. During this training session, participants performed 200 trials of one sequence (“trained”), 40 trials of a second sequence (“novice”), and 40 trials of un-cued pseudorandom sequences (“random”). The identity of each of the two sequences was selected from a set of three different sequences for each participant (3-2-4-1-2-4-2-1, 2-1-4-2-3-4-1-3, and 4-3-1-4-2-3-1-2). These sequences were chosen to be free of trills (e.g. 1-2-1) and repeats (e.g. 1-1). The selection of these sequences was counterbalanced across participants such that the trained sequence for one third of the subjects was the novice sequence in another third of the subjects and omitted for the final third of the subjects. This procedure ensured that our results would not be driven by the idiosyncrasies of the specific sequences chosen. In session two, participants performed 8 blocks of 30 trials, 120 trials for each sequence trained in session one. We imposed movement time limits on performance during session two, defined as the 70^th^ percentile of movement times in a warm-up session (see Figure S4 for plots of time limits for each subject and sequence during the test session). This warm-up session consisted of 60 trials of the trained sequence and 30 trials of the novice sequence performed in the scanner prior to the functional scans with the order of presentation randomized. Separate time limits were assigned for each sequence and each participant with a maximum of 3 s, and the time limit was reset to be stricter each block if participants’ accuracy was above 70% on the previous block. Each trial was associated with one of three incentive magnitudes: $5, $10, or $30. These reward values were presented simultaneously with the sequence color cues. The order of reward values was set to be an m-sequence to mitigate carryover effects (Buracas and Boynton 2002). Participants were informed that a trial would be selected at random at the end of the experiment, and that they would earn the associated reward provided they performed the target sequence without error under the time limit. Our metric of successful task performance therefore incorporated both speed and accuracy.

### Incentive-behavior analysis

We estimated the effect of reward on behavioral performance using a Bayesian hierarchical logistic regression model. The dependent variable was trial success. The model included fixed effects of $10-$5 and $30-$5 and allowed intercepts to vary by subject. We assigned weakly-informative N(0,1) priors to all parameters after standardizing the inputs (Gelman et al. 2008). The model was implemented using the R package {brms} (Bürkner 2017), which translates R-style regression formulae to code in the probabilistic programming language, Stan (Carpenter et al. 2017). Posterior parameter estimates were obtained using MCMC sampling (4 chains, 10000 iterations per chain, 50% warmup).

All MCMC chains passed visual inspection, all *Ȓ* values were 1, and all bulk and tail effective sample sizes (ESS) were large. After fitting the models, we performed graphical posterior predictive checks using the R packages {bayesplot} (Gabry et al. 2019) and {loo} (Vehtari et al. 2017). To quantify uncertainty about the effects of interest, we computed 95% highest density intervals (HDI) as well as probabilities of direction *(pd).* The HDI is defined such that, e.g., 95% of the distribution lies within the interval and every point inside the interval has higher credibility than every point outside the interval (Kruschke 2011). The *pd* is defined as the probability that an effect goes in the direction indicated by the median estimate (Makowski et al. 2019).

### fMRI acquisition and pre-processing

MR data were acquired with a 3T GE scanner (MR 750) with a 32-channel head coil. Functional data were obtained using a 1-shot multi-band T2*-weighted echo-planar imaging (EPI) sequence sensitive to blood oxygenation level-dependent (BOLD) contrast (TR = 1200 msec, TE = 30 msec, flip angle = 70°, 21 cm field of view, in-plane resolution = 2.4 mm × 2.4 mm, MB acceleration = 3). Each functional volume contained 51 contiguous 2.5 mm-thick axial slices separated by a 0 mm inter-slice gap acquired in an interleaved manner. Whole-brain T1-weighted scans were acquired for anatomical localization. Functional data were realigned to the third volume acquired, slice-time corrected, and registered to the MNI-152 template using a nonlinear warp. Functional data were smoothed with a 6mm FWHM kernel for the whole-brain univariate analysis but left unsmoothed for all multivariate analyses (see below). Pre-processing was performed using AFNI (Cox,1996).

### Whole-brain Univariate Analysis

We modeled BOLD responses to the preparatory cue using generalized linear models (GLM) implemented in using AFNI. As shown in Figure 1, the preparatory cue conveyed information about the reward that could be obtained (e.g., “$30”) and the sequence that should be performed (e.g., a blue square for sequence A). One GLM was used to capture the spatially distributed patterns of brain responses to the cue on each trial. This GLM modeled each trial’s response to the cue as a separate regressor, enabling us to obtain separate coefficient maps for each trial. The resulting coefficient maps were used as samples in a multivariate decoding analysis. This GLM also contained motor execution regressors for each sequence, which were created by convolving a gamma function with a square wave starting at movement onset with duration equal to the movement time. Another GLM was used to measure the extent to which BOLD responses were greater for larger monetary incentives. This GLM used the same motor execution regressors but contained 2 separate regressors for each sequence for the cue period. One regressor captured the mean response to the cue, while the other captured parametric modulation due to reward magnitude. This latter regressor was coded to predict linear changes in activity with reward magnitude while accounting for the mean response to the cue (with reward levels coded as 1,2, 3). The resulting coefficient maps (one per participant) were fed into a second-level group analysis using AFNI’s 3dttest++ command, which helped us identify the voxels whose activity reliably scaled with reward magnitude across participants.

To examine whether our multivariate analyses (see below) could be driven by mere univariate differences, we conducted additional second-level control analyses that examined several contrasts: 1) mean response of trained vs novice sequences, 2) linear effect of reward for trained vs novice sequences, 3) mean difference between correct and incorrect trials for trained vs novice sequences. For all second-level analyses, multiple comparisons corrections were performed via permutation testing with AFNI’s equitable thresholding and clustering (ETAC) procedure.

### Multivariate Analysis of reward-responsive regions

We used multivariate pattern analysis (MVPA) to localize the areas in the brain that contained information about the identity of an upcoming action and whether the action would be performed successfully (‘performance decoding’). Multivariate techniques—such as machine learning classifiers—are powerful tools for measuring information in the brain, because they are capable of discovering complex, high-dimensional mappings between spatially distributed patterns of brain activity and stimuli (or behaviors, or many other things the brain might contain information about). We performed MVPA using SpaceNet classifiers from the nilearn python package (Abraham et al., 2014), which use spatial priors and regularization to construct sparse but structured coefficient maps. When the classifier achieves high out-of-sample classification accuracy, it’s coefficient maps can be interpreted as information maps specifying voxels whose *joint activity* contains information about the class (Baldassarre et al. 2012; Michel et al. 2012; Grosenick et al. 2013; Dohmatob et al. 2014).

The classifiers were run separately on each subject and were given the trial-wise cue-related activity beta maps as input. We performed initial feature selection by considering only those voxels whose activity increased with reward magnitude at cue (group-level p < 0.001 uncorrected). We used a leave-one-participant-out approach to construct the masks to ensure a participant’s mask was independent from their samples (Esterman et al., 2010, Lee & Grafton, 2015). This masking reduced the dimensionality of the decoding problem and ensured that the decoders only used voxels that were modulated by reward. We ran one set of SpaceNets to classify patterns according to the sequence that was cued (‘action decoding’, Figure 4) and another set of SpaceNet classifiers which predicted whether the forthcoming trial would be performed successfully (‘performance decoding’, Figure 5). The SpaceNet classification procedure used an 80-20 train-test split with stratification to equalize the distribution of class labels across folds. We fed the resulting SpaceNet coefficient maps into a second-level conjunction analysis to produce a mask of voxels that were informative about both action and performance at the group level (Figure 6). Our inferences for the group-level maps assumed a cluster-corrected alpha threshold of p < 0.05. A bootstrap analysis confirmed that group-level mean classification accuracy was above 0.55 for action decoding and above 0.75 for performance decoding (Figure S5).

### Follow-up ROI Analysis

Next, we took a closer look at several anatomical sub-components of the action and performance information maps. After conjoining the action and performance information maps, we intersected the conjunction map with specific anatomical regions of interest, defined using the multi-modal Glasser atlas (Glasser et al. 2016). We focused here on the superior parietal lobule (parcel indices: 29, 45, 46, 47, 48, 49, 50), lateral prefrontal cortex (parcel indices: 67, 70, 71, 73, 79, 80, 81, 83, 84, 86), supplementary motor area (43, 44), pre-supplementary motor area (parcel index: 63), dorsal premotor cortex (parcel indices: 54, 55, 96, 97, 98), and primary motor cortex (parcel index: 8). For each of these ROIs, we considered only the right hemisphere, contralateral to the effector. For each subject, we performed 1001 permutations of a leave-one-trial-out cross-validated MVPA using a linear support vector classifier with default parameters in Scikit-learn (Pedregosa et al. 2012). In the first permutation, the classifiers were trained on data with true labels, but for the 1000 other permutations the labels for the training data were randomly shuffled. The classification target for the classifiers was action identity and the analysis yielded binary accuracy scores for each trial, representing whether the action cued on a trial was successfully decoded. With trial-wise decoding accuracies, we were able to test whether action decoding was enhanced for trials with larger incentives ($30) compared to trials with smaller incentives ($5, $10). We use permutation testing to assess whether decoding accuracy at each reward level was above chance and whether the differences in decoding accuracy between reward levels were greater than would be expected by chance. We compute p-values for these means using the formula (C + 1) / (N + 1) where N is the total number of permutations (1000) and C is the number of permutations whose means were greater than or equal to the ‘true’ mean. To address issues of multiple comparisons, we use 1000 bootstrap samples from the null distributions to determine the probability of obtaining statistically significant effects in zero, one, two, three, etc. ROIs under the null. This analysis showed that it was unlikely (p < .05) under the null to observe significant effects in multiple ROIs (Fig. S6).

### Brain-behavior Analysis

Lastly, we examined the link between action decoding and behavior. We used a hierarchical logistic regression model with Bayesian parameter estimation to model our trial success. This model was designed to test whether higher decoding accuracy in our ROIs was associated with higher behavioral accuracy. The dependent variable was behavioral performance (success or failure). The models included fixed effects of decoding accuracy

(correct or incorrect) and sequence identity (A or B) and allowed intercepts to vary by subject. The sequence identity predictor was included to control for the effect of sequence and thereby separate this effect from the effect of decoding accuracy.

### Data-availability

Data and code used in this project are available at https://github.com/adkinsty/dsp_scanner.

## Results

### Prospective reward improves motor sequence performance

Our participants were more likely to successfully complete sequences on $30 trials compared to both $10 trials (β_30-10_ = 0.14, CI_95%_ = [−0.04,0.29], pd = 94.5%; Fig. 2A) and $5 trials (β_30-5_ = 0.26, CI_95%_ = [0.09,0.42], pd = 99.9%). We imposed time limits on performance to match success rate across the two sequences (see Methods). However, we found evidence that accuracy was lower for the trained sequence compared to the novice sequence (β_T-N_ = −0.34, CI = [−0.44, −0.24], pd = 100%), suggesting that the time limits for the well-trained sequences were too strict. However, movement speed was greater for the trained than the novice sequence (β_T-N_ = 0.05, CI = [0.02,0.09], pd = 100%) and greater for $30 compared to $10 trials (β_30-10_ = 0.04, CI = [0.01,0.08], pd = 100%). We found little evidence of an interaction between reward size and sequence suggesting that performance on both sequences was similarly improved for larger rewards (β_30-10×T-N_ = 0.12, CI = [−0.12,0.35], pd = 84.7%; β_30-5×T-N_ = 0.04,CI = [−0.18,0.29], pd = 64.1%). In sum, our participants performed motor sequences more quickly and accurately when they were offered performance-contingent prospective reward of increasing size. In what follows, explore the hypothesis that this motivational enhancement of behavioral performance is related to motivational enhancements of the neural representations of action in the frontal cortex.

**Figure 2.**
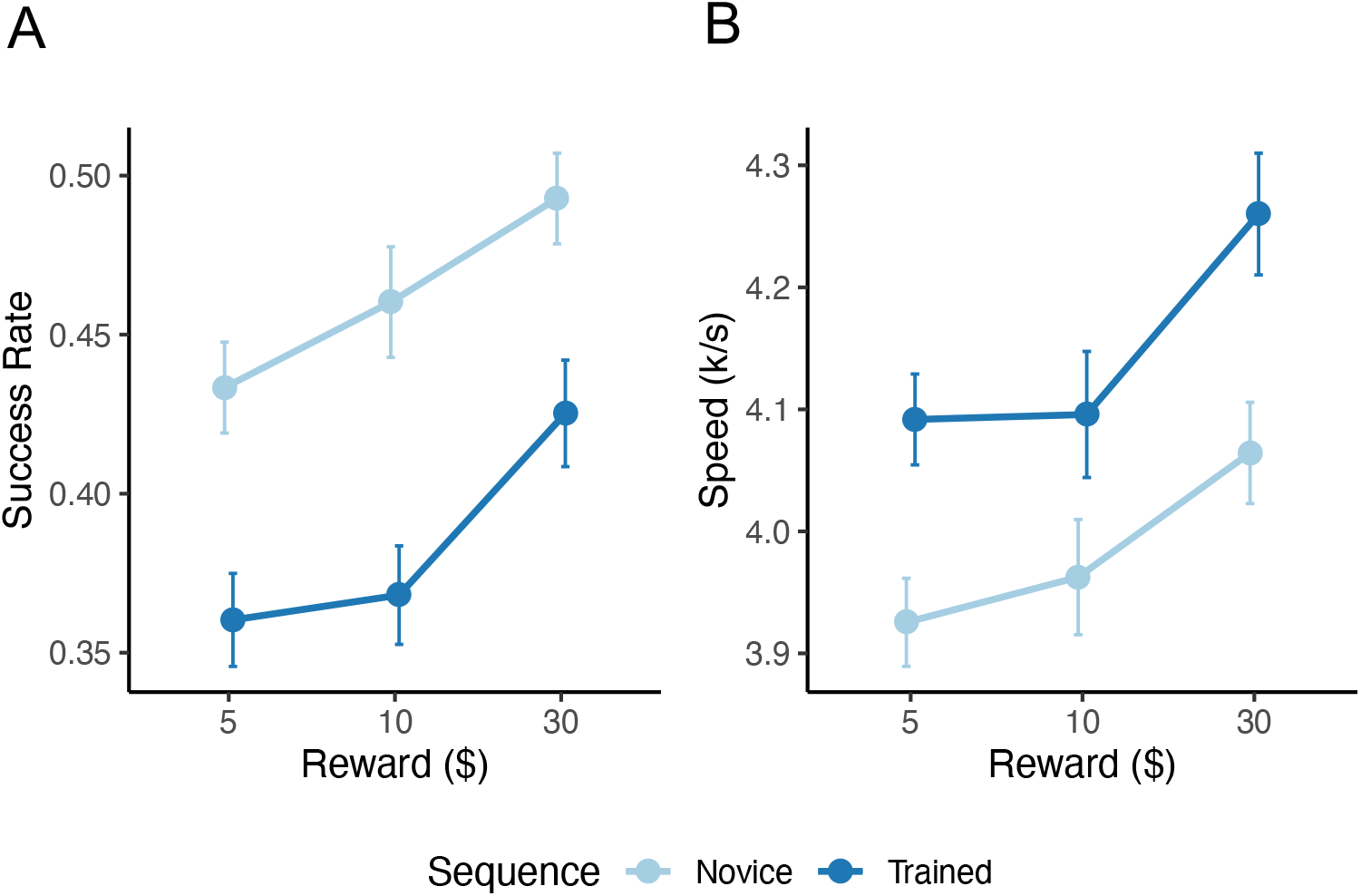
The effects of prospective reward size on subsequent behavioral performance. **A.** Participants were more likely to complete sequences under the prescribed time limits with increasing reward size. **B.** Participants pressed more keys per second with increasing reward size. Points reflect condition means and error bars reflect within-subject standard errors, calculated using the method described in (Cousineau 2005).

### Prospective reward increases preparatory activity in brain areas involved in motor planning

We first asked which regions of the brain were linearly responsive to motivation (i.e., prospective reward value) just prior to movement. This analysis revealed clusters of reward-related activity in most of the brain regions that we expected to be engaged during the DSP task, including LPFC, supplementary motor area, (SMA), pre-supplementary motor area (pre-SMA), dorsal premotor cortex (PMd), and the primary motor cortex (M1) (Figure 3, Table S1). Unsurprisingly, we also observed clusters of reward-related activity in reward regions such as the striatum and the pallidum. This analysis provided a reliable map of the brain regions whose activity was modulated by prospective rewards. We did not find any regions in which the extent of this reward modulation significantly differed between the sequences (whole-brain paired t-test, all P > 0.05, corrected). Of particular interest to this study is the finding that many regions involved in motor-skill performance were responsive to reward, including the primary motor cortex, premotor cortex and prefrontal cortex.

**Figure 3.**
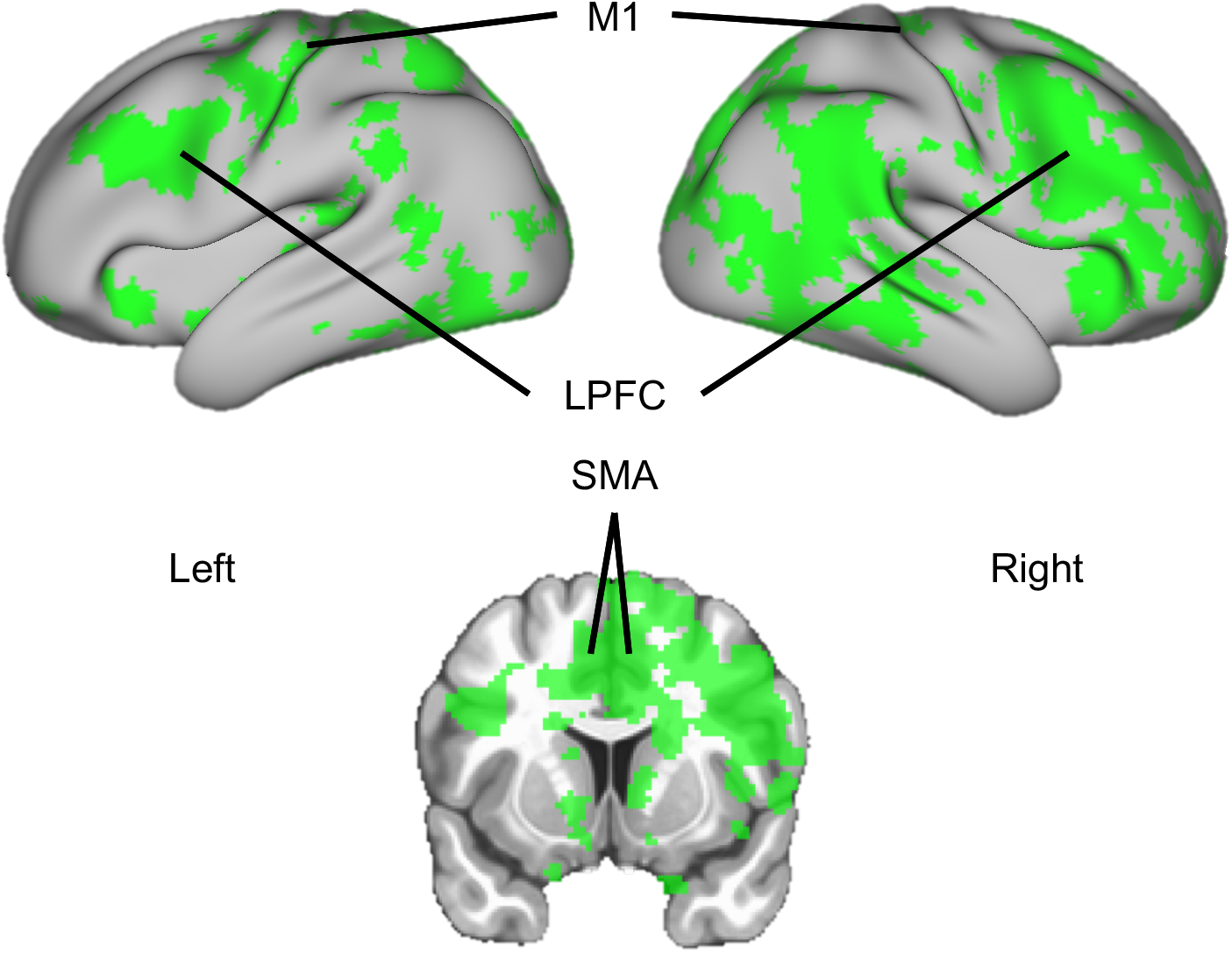
Reward-responsive brain areas. We used GLMs to test for effects of reward on hemodynamic responses to the pre-movement cue. The images above show the results of a group-level analysis of subject-level coefficient maps. For this visualization, we used threshold-free cluster correction and projected the results to surface space.

### Reward-modulated brain areas encode motor skill information

We performed MVPA using SpaceNet classifiers to identify the subset of regions from the reward map (Figure 3) whose patterns of activity also contained information about the upcoming action. Our first set of SpaceNet classifiers were trained to predict the identity of intended actions from patterns of brain activity preceding movement. At the group level, these classifiers predicted the intended action (for a held-out 20% of trials) with a mean accuracy of 0.61 and a standard deviation of 0.12 across subjects (P < .01; Figure 4A, Figure S5). We found action information distributed across the cortex including LPFC, SMA, and M1 (Figure 4B, Table S2). Importantly, this analysis considered only voxels that were responsive to reward magnitude. The regions in Figure 4B therefore simultaneously respond to changes in motivation and contribute to a representation of the action being prepared (Fig. S7). These findings are consistent with prior work showing converging reward and action signals in prefrontal areas and motor areas (McNamee et al., 2015).

**Figure 4.**
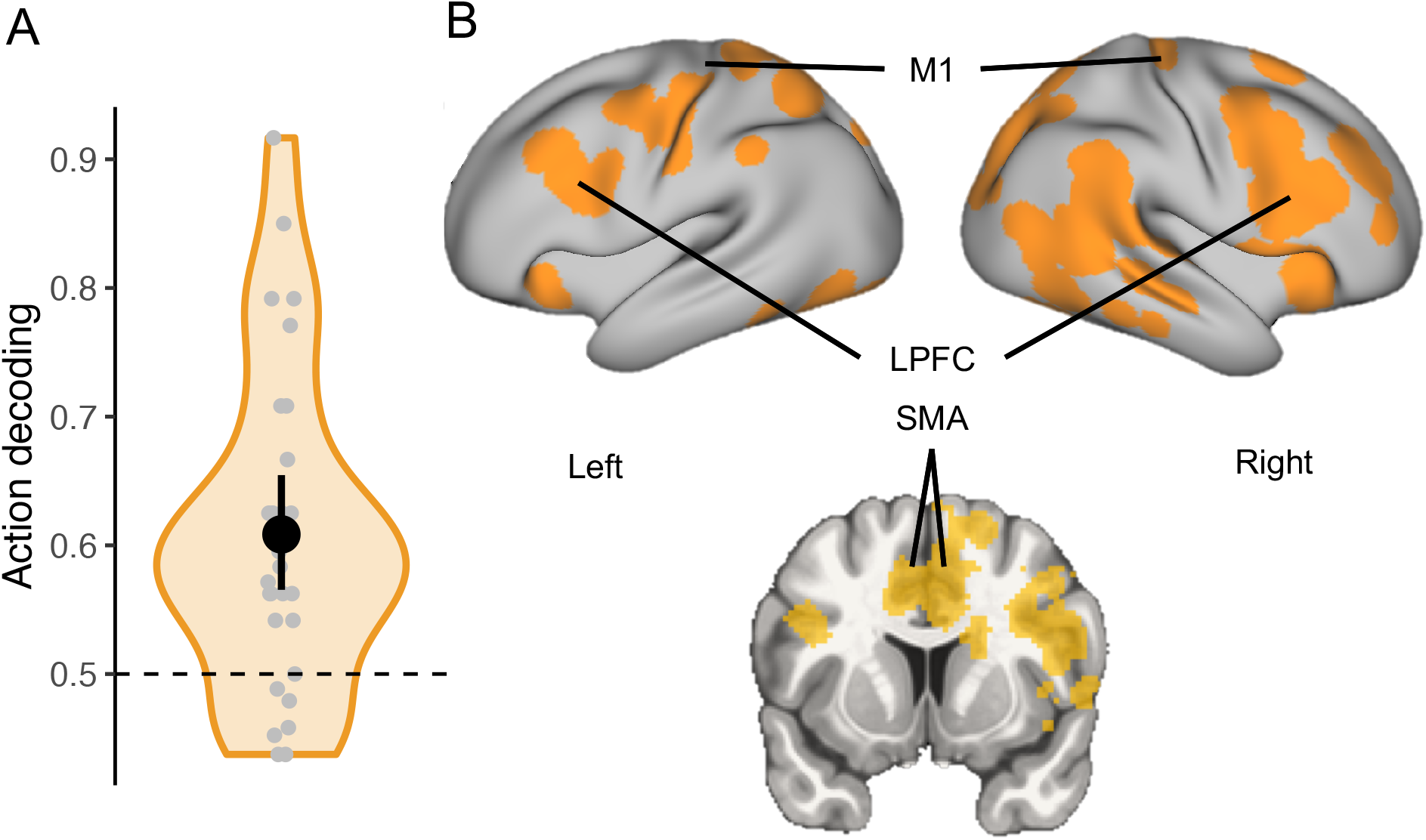
Action coding in reward-responsive brain areas. We trained multivariate classifiers to decode future action from patterns of hemodynamic responses to the pre-movement cue. This analysis only considered voxels that were responsive to reward (see Figure 3). (a) Violin plot of mean SpaceNet decoding accuracies for each subject. (b) Results of a group-level analysis of subject-level SpaceNet coefficient maps. For this visualization, we used threshold-free cluster correction, spatial smoothing with a 2mm kernel, and projected the results to surface space.

### Reward-modulated brain areas encode information about future performance

While information about reward and action appear to converge in a wide range of areas including LPFC, SMA, and M1, it remains a possibility that these areas have no practical relevance to future behavior. To test this, we ran a second set of classifiers that used patterns of activity prior to movement to predict whether a participant would be successful in the forthcoming DSP trial.

At the group level, the classifiers predicted future behavioral performance (on a held-out 20% of data) with a mean accuracy of 0.79 and standard deviation of 0.09 across subjects (p < .001, Figure 5A, Figure S5). This decoding analysis revealed a more restricted information map than the analysis of action decoding, but the map still included informative clusters in LPFC, SMA, and M1 (Figure 5B, Table S3).

**Figure 5.**
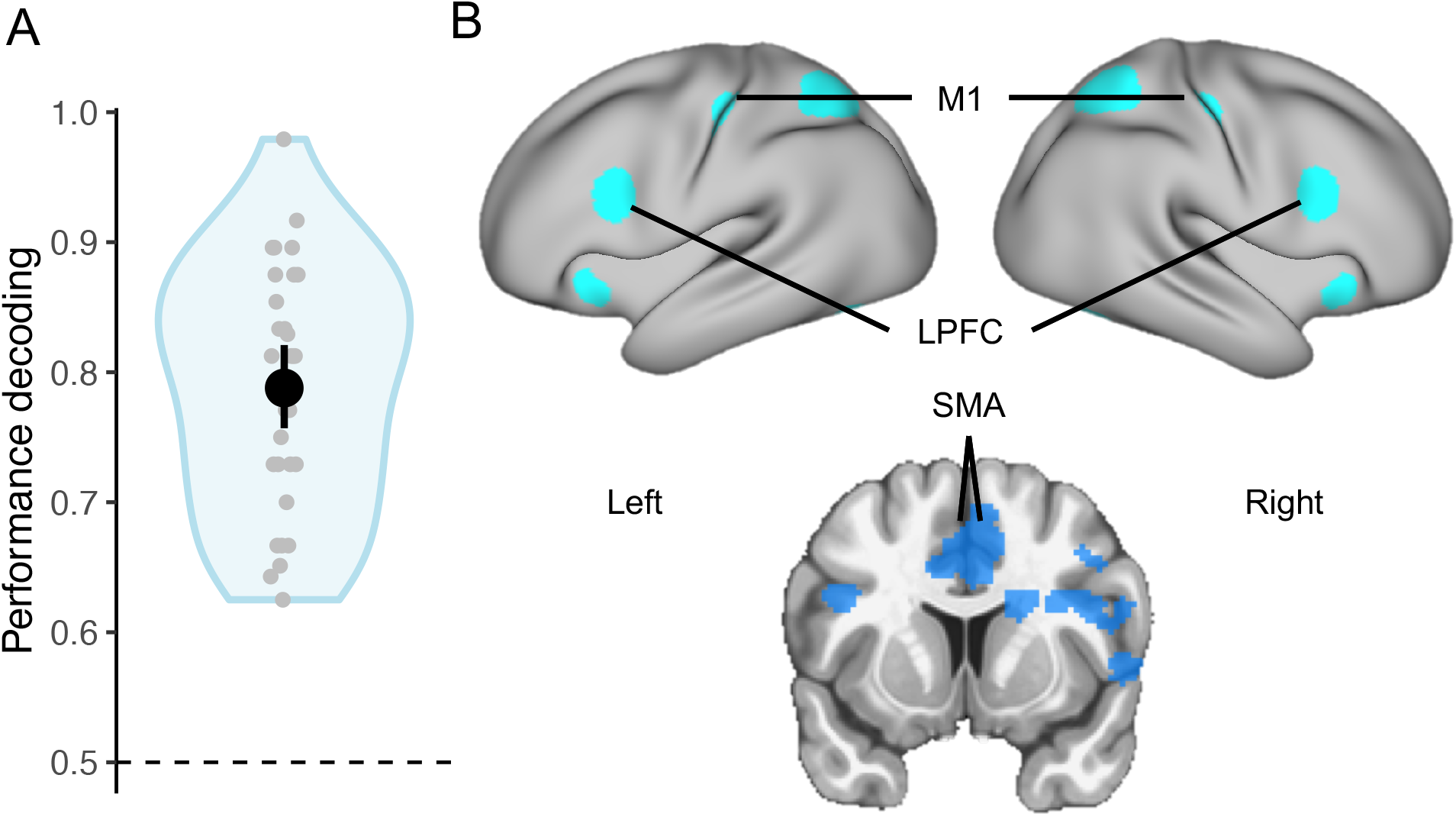
Performance-prediction in reward-responsive brain areas. We trained multivariate classifiers to predict future behavioral performance from patterns of hemodynamic responses to the pre-movement cue. This analysis only considered voxels that were responsive to reward (see Figure 3). (a) Violin plot of mean decoding accuracies for each subject. (b) Results of a group-level analysis of subject-level SpaceNet coefficient maps. For this visualization, we used threshold-free cluster correction, spatial smoothing with a 2mm kernel, and projected the results to surface space.

We performed several control analyses to ensure our classifiers were not merely detecting univariate differences between our conditions of interest (See Methods for a list of contrasts). These analyses did not reveal any significant clusters of activity when directly contrasting the mean univariate response between two sequences that survived multiple comparisons correction (no significant clusters of activity at p < 0.05). This suggests that our multivariate analyses were more sensitive in distinguishing between the two actions. Additionally, the linear effect of reward did not differ between the two sequences (no significant clusters of activity at p < 0.05). In another univariate control analysis, we tested whether the mean difference in cue-related activity for subsequently correct vs incorrect trials differed between the two sequences. We again found no significant clusters of activity in this analysis suggesting that the univariate response on both correct and incorrect trials was similar across the two sequences. Thus, although behavioral performance differs slightly between the sequences and at the different levels of reward, it is unlikely that our decoding results are driven by mere univariate differences.

Our fMRI results suggest that the reward-related regions identified previously are not only responsive to changes in motivation, but also relevant to future behavior. Furthermore, the information maps from our two MVPAs (Figures 4 and 5) were found to overlap in key regions of interest such as LPFC, SMA, and M1 (Table S4). This conjunction map, shown in Figure 6, delineates voxels that (1) respond to changes in motivation, (2) encode information about which action is being prepared, and (2) encode information about whether performance will be successful. We used this conjunctive information map as a mask for our subsequent ROI analyses.

**Figure 6.**
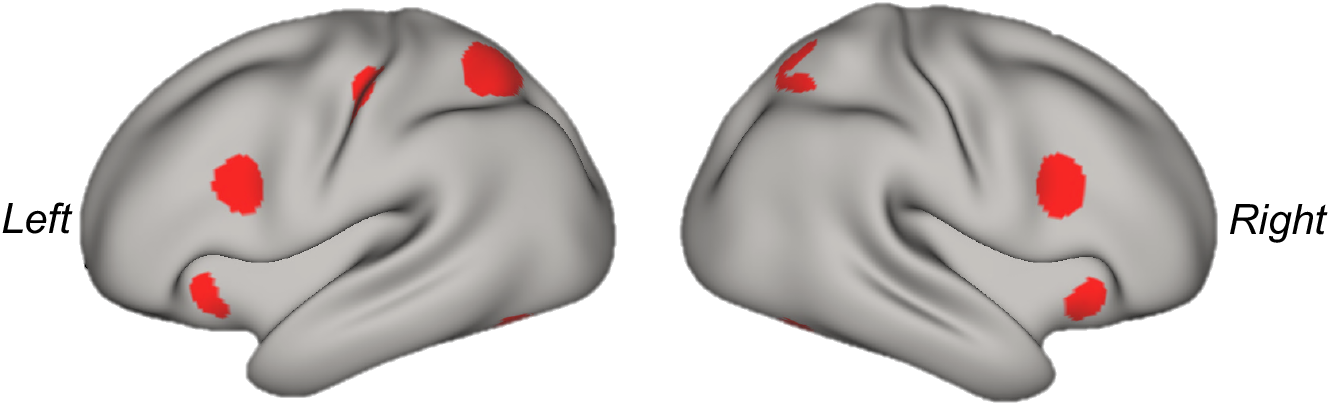
Overlap of action and performance information maps. The above mask includes voxels that were significant at p < 0.05 cluster-corrected in a group-level level conjunction analysis of the action and performance SpaceNet coefficient maps.

### Reward enhances action representations in frontal cortex

We hypothesized that motivation preferentially enhances cortical representations of skilled actions. While our multivariate analysis above revealed a widely distributed cortical representation of action that *jointly* carried information about skilled action, we were also interested whether skill representations can be identified in separate regions of interest previously implicated in studies of reward, action, and motor skills (Ballard et al. 2011; Wiestler and Diedrichsen 2013; Ramakrishnan et al. 2017; Yokoi et al. 2018; Yokoi and Diedrichsen 2019). To this end, we used linear support vector machines (SVM) to decode action identity from patterns of cue-related activity in the following ROIs in the right hemisphere (contralateral to the effector): LPFC, SMA, pre-SMA, PMd, and M1 (however, see Figure S2 for data from additional areas including the left hemisphere analogues to our ROIs). The purpose of this ROI analysis was to determine whether action information could be decoded locally in these ROIs and whether reward influenced this decoding.

Action decoding in pre-SMA was above chance for all reward levels ($5: *M* = .52, *P* = .03; $10: *M* = .52, *P* = .04; $30: *M* = .54, *P* < .001 and decoding accuracy was higher on $30 trials compared to $5 trials (*M*_diff_ = .02, *P* = .04). These results suggest that pre-SMA activity in isolation encodes information about action and that this activity may be most informative on high reward trials. For SMA, we found strong evidence that decoding was above chance for $30 trials (*M* = .52, *P* = .01) but not for $5 trials (*M* = .51, *P* = .09) or $10 trials (*M* = .51, *P* = .15). We observe a similar pattern of results for DLPFC ($5: *M* = .51, *P* = .18; $10: *M* = .5, *P* = .41; $30: *M* = .52, *P* = 0.06). A control analysis showed that the probability under the null of obtaining these consistent above chance classification on $30 trials across multiple regions was low (p < .05, Fig. S6). In fact, we obtained similar results in the left hemisphere ipsilateral to the hand used during performance (see Fig. S2). Interestingly, we could not decode upcoming actions above chance prior to movement from PMd (all p > .05). So, while this region may contribute to a whole-brain distributed representation of action, it does not in isolation carry enough information about action to support accurate decoding prior to movement (see Supplemental Figures S2A-E for the null distributions estimated using 1000 permutations of the classification procedure). Overall, these results suggest anterior control regions—DLPFC, SMA, and pre-SMA—locally encode upcoming actions during movement preparation specifically on high reward trials. Furthermore, in pre-SMA information about action appeared to increase with reward, suggesting that reward sharpens representations of action in pre-SMA during movement planning.

### Successful action decoding in SMA coincides with more accurate behavioral performance

Our results above suggest that motivation by prospective reward enhances action as well as neural representations of those actions in the brain. While these motivational effects may be coincidental, we considered the possibility that the enhanced action coding may be a neural mechanism by which subsequent behavior is enhanced. It follows from this hypothesis that behavioral performance should be better on trials in which action codes had high fidelity (i.e., when action identity was correctly decoded from preparatory brain activity).

We found evidence that participants were more likely to succeed when action could be decoded from preparatory activity in SMA (β_decode_ = 0.09, CI_95%_ = [-0.01,0.19], pd = 96.3%; Fig. 7B), however this effect was not significant for the other ROIs after controlling for the effect of sequence on behavioral performance (pd < 90%). The above results suggest that the motivational enhancement of action codes used in planning may be a key neuronal mechanism underlying the motivational enhancement of performance.

**Figure 7.**
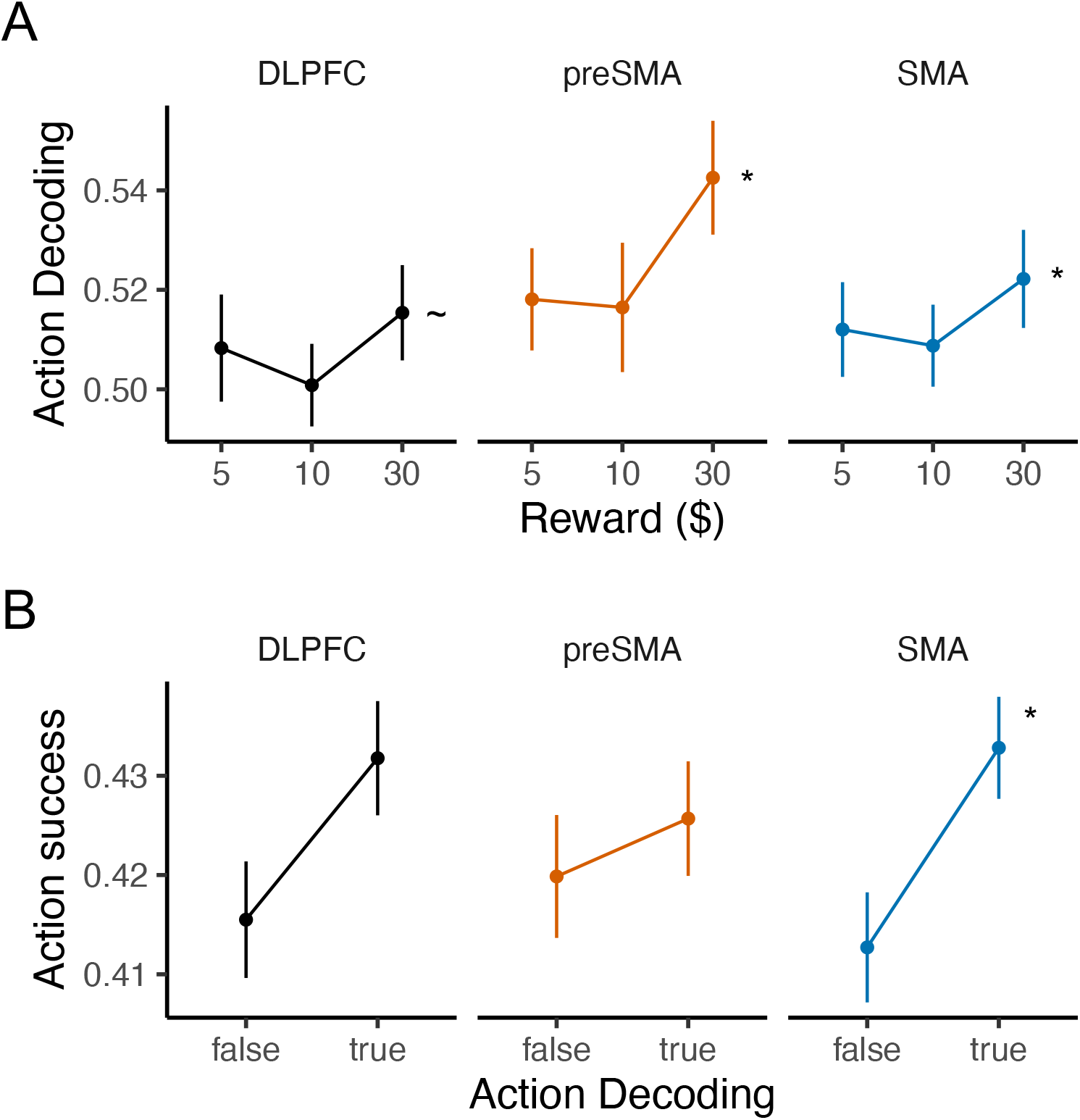
Effects of reward on action decoding and action decoding on performance. (A) We found evidence that action decoding from LPFC, pre-SMA, SMA was enhanced by the prospect of a $30 reward. Here we show group-mean decoding accuracies. The asterisks denote that decoding for $30 trials was significant at p < 0.05 and the tilde denotes that decoding was nearly significant at p = 0.06. Significance tests were performed at the group-level using 1000 random permutations of the classification procedure (i.e., by training on shuffled data). (B) We found evidence that action decoding in SMA was associated with better future behavioral performance, after controlling for action identity (i.e., sequence). The asterisk denotes pd > 95%. See Supplementary figures S3A and S3B for subject-level data.

## Discussion

In this study, we sought to examine the neural mechanisms that contribute to the motivational enhancement of action. We found that performance in a motor sequencing task improved as the size of prospective performance-contingent reward increased. When examining cue-related activity just before movement onset, we uncovered distributed patterns of activity across a large network of regions important for motor planning that simultaneously coded for reward value, action, and future behavioral success. We then interrogated a subset of these regions to examine how action coding was impacted by increasing reward values. We found that our ability to decode upcoming actions from pre-SMA improved as reward values increased. A follow-up analysis showed that people were more likely to succeed on trials in which we could correctly decode the upcoming action from preparatory activity in SMA. Our results suggest that incentive-motivated performance may depend on enhanced representations of action used in movement planning

### Reward modulates a widespread task-relevant network

Our results show that motivation (i.e, prospective reward value) modulated a widespread task network prior to movement (Figure 3). In these regions, the amplitude of the hemodynamic response to the cue was larger for high-value trials compared to low-value trials. This motivation-modulated network included canonical reward regions such as the striatum (Apicella et al. 1991; Samejima et al. 2005; Balleine et al. 2007; Delgado 2007) and ventromedial frontal cortex. However, we also found that prospective reward magnitude parametrically increased activity in many regions thought to be involved in motor planning including lateral frontal cortex and supplementary motor areas. This finding is consistent with prior work showing anticipatory reward modulation in task-relevant brain areas in cognitive and motor tasks (Leon and Shadlen 1999; Knutson et al. 2003; Roesch and Olson 2003, 2004; Wallis and Miller 2003; Kennerley and Wallis 2009; Wallis and Kennerley 2010; Peterson and Seger 2013; Marsh et al. 2015; Ramkumar et al. 2016; Ramakrishnan et al. 2017; Galaro et al. 2018)

### Multiplexed code for reward, action, and performance

To investigate how prospective reward relates to the neural coding of action, we sought to determine which of these reward-responsive areas specifically carried information about both the identity and the quality of the upcoming action. Using multivariate decoding, we found that distributed patterns of activity across the cortex simultaneously carried information about prospective reward value, action and performance. Previous studies have shown that reward signals and action signals converge in several areas including the prefrontal cortex (Kargo et al. 2007; FitzGerald et al. 2012; Cai and Padoa-Schioppa 2014; Hunt et al. 2015; McNamee et al. 2015), the striatum (Kargo et al. 2007; FitzGerald et al. 2012; Cai and Padoa-Schioppa 2014; Hunt et al. 2015; McNamee et al. 2015), anterior cingulate cortex (Hayden and Platt 2010), and primary motor cortex (Ramakrishnan et al. 2017; Galaro et al. 2018). We similarly find that information about reward, action and future performance converge in lateral prefrontal cortex, posterior parietal cortex, and more dorsal regions of medial frontal cortex (pre-SMA and SMA).

Explicit cueing of a sequential action as employed in this study enables advanced motor planning (Diedrichsen and Kornysheva 2015). Especially early on in training, cognitive control processes have been shown to aid in the performance of motor sequencing skills. For example, working memory capacity relates to the rate of skill learning (Anguera et al. 2011) and novice motor sequence performance coincides with increased activity in LPFC (Grafton et al. 1995; Willingham et al. 2002; Schendan et al. 2003). It is thought that information about prospective rewards is introduced to goal-directed action networks through LPFC (Ballard et al. 2011). It has been demonstrated that performance-contingent rewards can enhance cognitive control. Rewards have been associated with improvements in the advanced maintenance of goal information (Jimura et al. 2010; Botvinick and Braver 2014; Etzel et al. 2016) and research has demonstrated motivational enhancements of representations in PFC (Leon and Shadlen 1999; Kobayashi et al. 2002; Watanabe et al. 2002; Kargo et al. 2007; Etzel et al. 2016; Parro et al. 2018). We therefore reason that prospective rewards may induce a preferential enhancement of task-relevant representations—that is, representations used in the planning of future performance (Botvinick and Braver 2014; Westbrook and Braver 2015; Yee and Braver 2018).

### Enhanced action representations as a mechanism for enhanced performance

Our multivariate analyses showed that motivational signals and preparatory action representations converge in several brain areas. While such convergence is likely necessary for motivation to enhance behavior, it is unclear exactly what happens when these signals converge. One possibility is that motivation enhances cortical representations of upcoming actions. We addressed this possibility in follow-up ROI analyses and found evidence for such enhancement in pre-SMA (Figure 7A). Our results also show that anticipatory action decoding in SMA was linked to subsequent behavioral performance (Figure 7B). Together, these results provide support for the hypothesis that the prospect of reward may lead to enhanced performance by enhancing prospective representations of action used in planning.

Although our a priori focus was on LPFC as a key area involved in the enhancement of action by reward, our results predominantly implicate SMA and pre-SMA. It is known that cells in the supplementary motor area encode information about the sequential order of future movements (Tanji & Shima, 1994), while cells in pre-SMA are responsible for updating such sequential movement plans (Shima et al., 1996). This was corroborated by more recent work demonstrating a deficit in the inhibition of planned movements following lesion to the right pre-SMA (Nachev et al., 2007). These findings suggest that the SMA and pre-SMA play critical roles in the planning of sequential actions. While it has been shown that the supplementary motor area encode information about expected reward (Campos et al., 2005), we provide evidence that reward modulates the fidelity of putative action plans.

### Limitations

It is possible that reward also enhances performance through enhanced processing during the execution period itself. However, we chose to focus on activity at cue (motor preparation) rather than at movement (motor execution) because it was less influenced by confounds. First, in the present experiment, the duration of the movement period varied considerably across time, action, and participant. Second, error processing during execution could contribute to the decoding of successful performance. Third, relatively strict time limits were used to prevent the strategic slowing of execution speed on high reward trials. This design choice led to a relatively high number of errors during execution, limiting the number of successfully completed trials that could be used for analysis of the movement period itself. On error trials, the performed sequence of button presses was be quite different within a sequence (i.e., fewer movements and erroneous movements) and added greatly to the variability in the patterns of BOLD activity during this period. Nonetheless, it is important for future work to consider the effects of motivation on brain activity during execution, especially because this may be the locus of enhancement for performance that depends less on anticipatory cognitive processing (e.g., the performance of expert motor skills).

We think it is plausible that the action decoding we observed was driven by action identity (e.g., sequence order) and that these representations became more distinct with increased motivation. However, we cannot rule out the possibility that our decoding analyses distinguished neuronal patterns based on skill-level rather than sequence per se. On this view, there may be functional neural correlates of novice and trained skills that do not depend much on the identity of the action, such as the motor sequence order. There is evidence that novice and expert skills depend on distinct neuronal mechanisms (Dayan and Cohen 2011; Yokoi and Diedrichsen 2019), and such differences could conceivably be detected by MVPA decoding techniques. However, none of the skills in our task were expert, having been trained for only a few hundred trials at most. If decoding were driven by skill-level, it would be curious why such brain differences in skill level would be enhanced by reward. One possibility is that novice and trained skills differentially engage higher cognitive processes (Poldrack et al. 2005). If reward modulates these processes, it may result in increased differentiation between patterns of activity associated with novice and trained skills. Regardless, it is clear that preparatory activity differs between the two sequences, that motivation enhances the distinctiveness of this preparatory activity, and that this increased distinctiveness coincides to better task performance.

Our analyses also focused heavily on decoding action information from patterns of cortical activity. We focused on this aspect of our dataset because prior work validated the approach of measuring cortical representations using MVPA (Wiestler and Diedrichsen 2013; Nambu et al. 2015; Yokoi et al. 2018; Yokoi and Diedrichsen 2019) and because recent work has shown that multivariate decoding from cortex can be affected by motivation (Etzel et al. 2016; Parro et al. 2018). However, motivated action depends on cortico-striatal loops (Doyon et al. 2003; Turner and Desmurget 2010; Dayan and Cohen 2011; Frank 2011; Dudman and Krakauer 2016; Hikosaka et al. 2018) and reward increases activity in the basal ganglia during motor learning (Wachter et al., 2009). A recent study showed that showed that training with reward in an SRT task was associated with increased functional connectivity between premotor cortex and the cerebellum as well as the premotor cortex and the striatum (Steel et al., 2019). Very few studies have used neuroimaging to address reward-motor interactions and we must note that those by Wächter, et al. and Steel, et al. both examined the impact of reward during the acquisition of motor sequencing skill rather than in motivating the execution of a previously learned skill as we do here. Nevertheless, future work should examine whether cortico-striatal or cortico-cerebellar loops are related to the motivational enhancements of performance and neural decoding reported in the present work.

### Closing remarks

In sum, we provide evidence that behavioral performance is enhanced by motivation and that a widespread network of motor planning regions jointly contains information about reward, action, and performance. Additionally, our ability to decode skilled actions from patterns of BOLD activity from these regions in isolation was enhanced by the prospect of large rewards. We propose that motivation increases the expected value of deploying attention to action representations used in planning, thereby increasing their signal to noise ratio and consequently improving behavioral performance.

## Notes

### Competing Interest Statement

The authors have declared no competing interest.

## Works Cited

Abraham, A., Pedregosa, F., Eickenberg, M., Gervais, P., Mueller, A., Kossaifi, J & Varoquaux, G. 2014. Machine learning for neuroimaging with scikit-learn. Frontiers in neuroinformatics, 8, 14

Anderson, S. P., Adkins, T. J., Gary, B. S., & Lee, T. G. 2020. Rewards interact with explicit knowledge to enhance skilled motor performance. Journal of Neurophysiology, 123:6, 2476–2490.

Anguera JA, Reuter-Lorenz PA, Willingham DT, Seidler RD. 2011. Failure to Engage Spatial Working Memory Contributes to Age-related Declines in Visuomotor Learning. Journal of Cognitive Neuroscience. 23:11–25.

Apicella P, Ljungberg T, Scarnati E, Schultz W. 1991. Responses to reward in monkey dorsal and ventral striatum. Experimental brain research. 85:491–500.

Baldassarre L, Mourão-Miranda J, Pontil M. 2012. Structured Sparsity Models for Brain Decoding from fMRI data. 2012 Second International Workshop on Pattern Recognition in NeuroImaging. 1:5–8.

Ballard IC, Murty VP, Carter RM, MacInnes JJ, Huettel SA, Adcock RA. 2011. Dorsolateral Prefrontal Cortex Drives Mesolimbic Dopaminergic Regions to Initiate Motivated Behavior. The Journal of Neuroscience. 31:10340–10346.

Balleine BW, Delgado MR, Hikosaka O. 2007. The Role of the Dorsal Striatum in Reward and Decision-Making. The Journal of Neuroscience. 27:8161–8165.

Botvinick M, Braver T. 2014. Motivation and cognitive control: from behavior to neural mechanism. Annual review of psychology. 66:83–113.

Buracas GT, Boynton GM. 2002. Efficient design of event-related fMRI experiments using M-sequences. NeuroImage. 16:801–813.

Bürkner P-C. 2017. brms: An R Package for Bayesian Multilevel Models Using Stan. Journal of Statistical Software. 80.

Cai X, Padoa-Schioppa C. 2014. Contributions of Orbitofrontal and Lateral Prefrontal Cortices to Economic Choice and the Good-to-Action Transformation. Neuron. 81:1140–1151.

Campos, M., Breznen, B., Bernheim, K., & Andersen, R. A. 2005. Supplementary motor area encodes reward expectancy in eye-movement tasks. Journal of Neurophysiology.

Carpenter B, Gelman A, Hoffman MD, Lee D, Goodrich B, Betancourt M, Brubaker M, Guo J, Li P, Riddell A. 2017. Stan: A Probabilistic Programming Language. Journal of Statistical Software. 76.

Cousineau D. 2005. Confidence intervals in within-subject designs: A simpler solution to Loftus and Masson’s method. Tutorials Quantitative Methods Psychology. 1:42–45.

Cox, R. W. 1996. AFNI: software for analysis and visualization of functional magnetic resonance neuroimages. Computers and Biomedical research, 29:3, 162-173.

Dayan E, Cohen LG. 2011. Neuroplasticity Subserving Motor Skill Learning. Neuron. 72:443–454.

Delgado MR. 2007. Reward-related responses in the human striatum. Annals of the New York Academy of Sciences. 1104:70–88.

Diedrichsen J, Kornysheva K. 2015. Motor skill learning between selection and execution. Trends in Cognitive Sciences. 19:227–233.

Dohmatob E, Gramfort A, Thirion B, Varoquaux G. 2014. Benchmarking solvers for TV-L1 leastsquares and logistic regression in brain imaging. 2014 International Workshop on Pattern Recognition in Neuroimaging.

Doyon J, Penhune V, Ungerleider LG. 2003. Distinct contribution of the cortico-striatal and cortico-cerebellar systems to motor skill learning. Neuropsychologia. 41:252–262.

Dudman JT, Krakauer JW. 2016. The basal ganglia: from motor commands to the control of vigor. Current Opinion in Neurobiology. 37:158–166.

Engelmann JB, Pessoa L. 2007. Motivation sharpens exogenous spatial attention. Emotion (Washington, DC). 7:668–674.

Esterman, M., Tamber-Rosenau, B. J., Chiu, Y. C., & Yantis, S. 2010. Avoiding nonindependence in fMRI data analysis: leave one subject out. Neuroimage, 50:2, 572-576.

Etzel JA, Cole MW, Zacks JM, Kay KN, Braver TS. 2016. Reward Motivation Enhances Task Coding in Frontoparietal Cortex. Cerebral Cortex. 26:1647–1659.

FitzGerald THB, Friston KJ, Dolan RJ. 2012. Action-Specific Value Signals in Reward-Related Regions of the Human Brain. The Journal of Neuroscience. 32:16417–16423.

Frank MJ. 2011. Computational models of motivated action selection in corticostriatal circuits. Current Opinion in Neurobiology. 21:381–386.

Gabry J, Simpson D, Vehtari A, Betancourt M, Gelman A. 2019. Visualization in Bayesian workflow. J Royal Statistical Soc Ser Statistics Soc. 182:389–402.

Galaro JK, Celnik P, Chib VS. 2018. Motor Cortex Excitability Reflects the Subjective Value of Reward and Mediates Its Effects on Incentive-Motivated Performance. The Journal of Neuroscience. 39:1236–1248.

Gelman A, Jakulin A, Pittau MG, Su Y-S. 2008. A weakly informative default prior distribution for logistic and other regression models. Ann Appl Statistics. 2:1360–1383.

Glasser MF, Coalson TS, Robinson EC, Hacker CD, Harwell J, Yacoub E, Ugurbil K, Andersson J, Beckmann CF, Jenkinson M, Smith SM, Essen DC. 2016. A multi-modal parcellation of human cerebral cortex. Nature. 536:171–178.

Grafton S, Hazeltine E, Ivry R. 1995. Functional mapping of sequence learning in normal humans. Journal of cognitive neuroscience. 7:497–510.

Grosenick L, Klingenberg B, Katovich K, Knutson B, Taylor JE. 2013. Interpretable brain-wide prediction analysis with GraphNet. NeuroImage. 72:304–321.

Hayden BY, Platt ML. 2010. Neurons in Anterior Cingulate Cortex Multiplex Information about Reward and Action. The Journal of Neuroscience. 30:3339–3346.

Hikosaka O, Ghazizadeh A, Griggs W, Amita H. 2018. Parallel basal ganglia circuits for decision making. Journal of Neural Transmission. 125:515–529.

Hikosaka, O., Nakamura, K., Sakai, K., & Nakahara, H. 2002. Central mechanisms of motor skill learning. Current opinion in neurobiology, 12:2, 217-222.

Hunt LT, Behrens TE, Hosokawa T, Wallis JD, Kennerley SW. 2015. Capturing the temporal evolution of choice across prefrontal cortex. eLife. 4:e11945.

Jimura K, Locke HS, Braver TS. 2010. Prefrontal cortex mediation of cognitive enhancement in rewarding motivational contexts. Proceedings of the National Academy of Sciences. 107:8871–8876.

Kargo WJ, Szatmary B, Nitz DA. 2007. Adaptation of Prefrontal Cortical Firing Patterns and Their Fidelity to Changes in Action-Reward Contingencies. The Journal of Neuroscience. 27:3548–3559.

Kennerley SW, Wallis JD. 2009. Reward-dependent modulation of working memory in lateral prefrontal cortex. The Journal of neuroscience: the official journal of the Society for Neuroscience. 29:3259–3270.

Knutson B, Fong GW, Bennett SM, Adams CM, Hommer D. 2003. A region of mesial prefrontal cortex tracks monetarily rewarding outcomes: characterization with rapid event-related fMRI. NeuroImage. 18:263–272.

Kobayashi S, Lauwereyns J, Koizumi M, Sakagami M, Hikosaka O. 2002. Influence of Reward Expectation on Visuospatial Processing in Macaque Lateral Prefrontal Cortex. Journal of Neurophysiology. 87:1488–1498.

Kool W, Botvinick M. 2018. Mental labour. Nature human behaviour. 2:899–908.

Krakauer JW, Hadjiosif AM, Xu J, Wong AL, Haith AM. 2019. Motor Learning. Comprehensive Physiology.

Krawczyk DC, Gazzaley A, D’Esposito M. 2007. Reward modulation of prefrontal and visual association cortex during an incentive working memory task. Brain research. 1141:168–177.

Kruschke JK. 2011. Bayesian Assessment of Null Values Via Parameter Estimation and Model Comparison. Perspectives on Psychological Science. 6:299–312.

Lee, T. G., & Grafton, S. T. 2015. Out of control: Diminished prefrontal activity coincides with impaired motor performance due to choking under pressure. NeuroImage, 105, 145–155.

Leon MI, Shadlen MN. 1999. Effect of Expected Reward Magnitude on the Response of Neurons in the Dorsolateral Prefrontal Cortex of the Macaque. Neuron. 24:415–425.

Makowski D, Ben-Shachar MS, Chen SHAH, Lüdecke D. 2019. Indices of Effect Existence and Significance in the Bayesian Framework. Frontiers in psychology. 10:2767.

Marsh BT, Tarigoppula VSA, Chen C, Francis JT. 2015. Toward an Autonomous Brain Machine Interface: Integrating Sensorimotor Reward Modulation and Reinforcement Learning. The Journal of Neuroscience. 35:7374–7387.

McNamee D, Liljeholm M, Zika O, O’Doherty JP. 2015. Characterizing the Associative Content of Brain Structures Involved in Habitual and Goal-Directed Actions in Humans: A Multivariate fMRI Study. The Journal of Neuroscience. 35:3764–3771.

Michel V, Gramfort A, Eger E, Varoquaux G, Thirion B. 2012. A Comparative Study of Algorithms for Intra-and Inter-subjects fMRI Decoding. springer.

Mosberger AC, Clauser L de, Kasper H, Schwab ME. 2016. Motivational state, reward value, and Pavlovian cues differentially affect skilled forelimb grasping in rats. Learning & memory (Cold Spring Harbor, NY). 23:289–302.

Nachev, P., Wydell, H., O’neill, K., Husain, M., & Kennard, C. 2007. The role of the pre-supplementary motor area in the control of action. Neuroimage, 36, T155–T163.

Nambu I, Hagura N, Hirose S, Wada Y, Kawato M, Naito E. 2015. Decoding sequential finger movements from preparatory activity in higher-order motor regions: a functional magnetic resonance imaging multi-voxel pattern analysis. European Journal of Neuroscience. 42:2851–2859.

Nissen M, Bullemer P. 1987. Attentional requirements of learning: Evidence from performance measures. Cognitive Psychology. 19:1–32.

Parro C, Dixon ML, Christoff K. 2018. The neural basis of motivational influences on cognitive control. Human Brain Mapping. 39:5097–5111.

Patzelt EH, Kool W, Millner AJ, Gershman SJ. 2018. Incentives Boost Model-Based Control Across a Range of Severity on Several Psychiatric Constructs. Biological Psychiatry. 85:425–433.

Pedregosa F, Varoquaux G, Gramfort A, Michel V, Thirion B, Grisel O, Blondel M, Müller A, Nothman J, Louppe G, Prettenhofer P, Weiss R, Dubourg V, Vanderplas J, Passos A, Cournapeau D, Brucher M, Perrot M, Duchesnay É. 2012. Scikit-learn: Machine Learning in Python. arXiv.

Peterson EJ, Seger CA. 2013. Many hats: intratrial and reward level-dependent BOLD activity in the striatum and premotor cortex. Journal of neurophysiology. 110:1689–1702.

Poldrack, R. A., Sabb, F. W., Foerde, K., Tom, S. M., Asarnow, R. F., Bookheimer, S. Y., & Knowlton, B. J. (2005). The neural correlates of motor skill automaticity. Journal of Neuroscience, 25(22), 5356–5364.

Ramakrishnan A, Byun YW, Rand K, Pedersen CE, Lebedev MA, Nicolelis MAL. 2017. Cortical neurons multiplex reward-related signals along with sensory and motor information. Proceedings of the National Academy of Sciences. 114:E4841–E4850.

Ramkumar P, Dekleva B, Cooler S, Miller L, Kording K. 2016. Premotor and Motor Cortices Encode Reward. PLOS ONE. 11:e0160851.

Roesch MR, Olson CR. 2003. Impact of Expected Reward on Neuronal Activity in Prefrontal Cortex, Frontal and Supplementary Eye Fields and Premotor Cortex. Journal of Neurophysiology. 90:1766–1789.

Roesch MR, Olson CR. 2004. Neuronal activity related to reward value and motivation in primate frontal cortex. Science (New York, NY). 304:307–310.

Samejima K, Ueda Y, Doya K, Kimura M. 2005. Representation of Action-Specific Reward Values in the Striatum. Science. 310:1337–1340.

Schendan HE, Searl MM, Melrose RJ, Stern CE. 2003. An fMRI Study of the Role of the Medial Temporal Lobe in Implicit and Explicit Sequence Learning. Neuron. 37:1013–1025.

Seidler RD, Bo J, Anguera JA. 2012. Neurocognitive Contributions to Motor Skill Learning: The Role of Working Memory. Journal of Motor Behavior. 44:445–453.

Serences JT. 2008. Value-based modulations in human visual cortex. Neuron. 60:1169–1181.

Turner RS, Desmurget M. 2010. Basal ganglia contributions to motor control: a vigorous tutor. Current Opinion in Neurobiology. 20:704–716.

Shima, K., Mushiake, H., Saito, N., & Tanji, J. 1996. Role for cells in the presupplementary motor area in updating motor plans. Proceedings of the National Academy of Sciences, 93:16, 8694-8698.

Steel, A., Silson, E. H., Stagg, C. J., & Baker, C. I. 2019. Differential impact of reward and punishment on functional connectivity after skill learning. NeuroImage, 189, 95–105.

Tanji, J., & Shima, K. 1994. Role for supplementary motor area cells in planning several movements ahead. Nature, 371:6496, 413-416.

Vehtari A, Gelman A, Gabry J. 2017. Practical Bayesian model evaluation using leave-one-out cross-validation and WAIC. Stat Comput. 27:1413–1432.

Wächter T, Lungu OV, Liu T, Willingham DT, Ashe J. 2009. Differential effect of reward and punishment on procedural learning. The Journal of neuroscience: the official journal of the Society for Neuroscience. 29:436–443.

Wallis JD, Kennerley SW. 2010. Heterogeneous reward signals in prefrontal cortex. Current opinion in neurobiology. 20:191–198.

Wallis JD, Miller EK. 2003. Neuronal activity in primate dorsolateral and orbital prefrontal cortex during performance of a reward preference task. European Journal of Neuroscience. 18:2069–2081.

Watanabe M, Hikosaka K, Sakagami M, Shirakawa S. 2002. Coding and monitoring of motivational context in the primate prefrontal cortex. The Journal of neuroscience: the official journal of the Society for Neuroscience. 22:2391–2400.

Westbrook A, Braver TS. 2015. Cognitive effort: A neuroeconomic approach. Cognitive, affective & behavioral neuroscience. 15:395–415.

Wiestler T, Diedrichsen J. 2013. Skill learning strengthens cortical representations of motor sequences. eLife. 2:e00801.

Willingham DB, Salidis J, Gabrieli JD. 2002. Direct Comparison of Neural Systems Mediating Conscious and Unconscious Skill Learning. Journal of Neurophysiology. 88:1451–1460.

Yee DM, Braver TS. 2018. Interactions of motivation and cognitive control. Current Opinion in Behavioral Sciences. 19:83–90.

Yokoi A, Arbuckle SA, Diedrichsen J. 2018. The Role of Human Primary Motor Cortex in the Production of Skilled Finger Sequences. Journal of Neuroscience. 38:1430–1442.

Yokoi A, Diedrichsen J. 2019. Neural Organization of Hierarchical Motor Sequence Representations in the Human Neocortex. Neuron. 103:1178–1190.e7.

