## Supplemental figures and tables for "Reward modulates cortical representations of action"

Supplementary Materials for
"Reward modulates cortical representations of action"

Tyler J. Adkins[1*], Taraz G. Lee[1]
Department of Psychology, University of Michigan, Ann Arbor, MI, 48109

[*]Corresponding author's email:


**Figure S1.** *Distributions of performance speed at the start and end of training for each sequence*. Speed was faster at the end of training for the trained sequences but not the random sequences. This result demonstrates sequence learning occurred in training. The large points are group-level means and the smaller points are subject-level means.

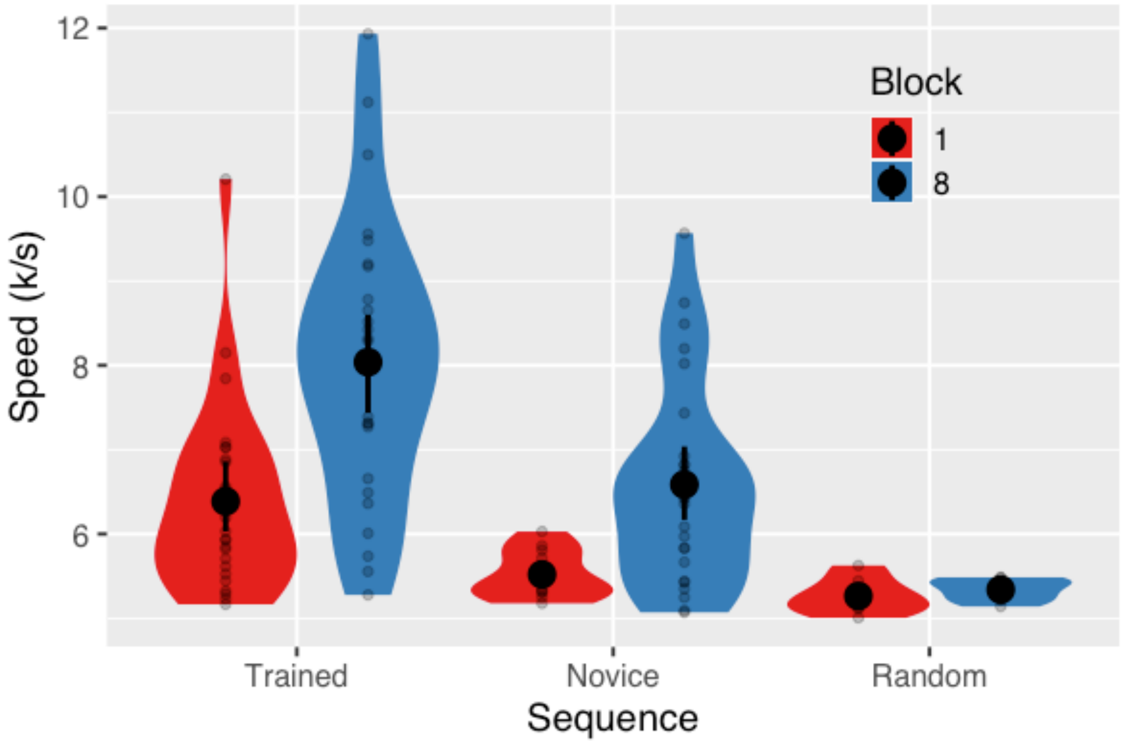

**Figure S2A.** *$5 ROI Decoding vs chance.* Each panel shows the distribution of action decoding accuracies for 1000 classifiers trained on shuffled data. The red vertical line represents the 95th percentile of the null distribution while the green vertical line represents the group mean decoding accuracy for classifiers trained on true data. If the green line exceeds the red, then $p < 0.05$ that the classification accuracy is at chance. In the panels below the L prefix denotes left hemisphere regions.

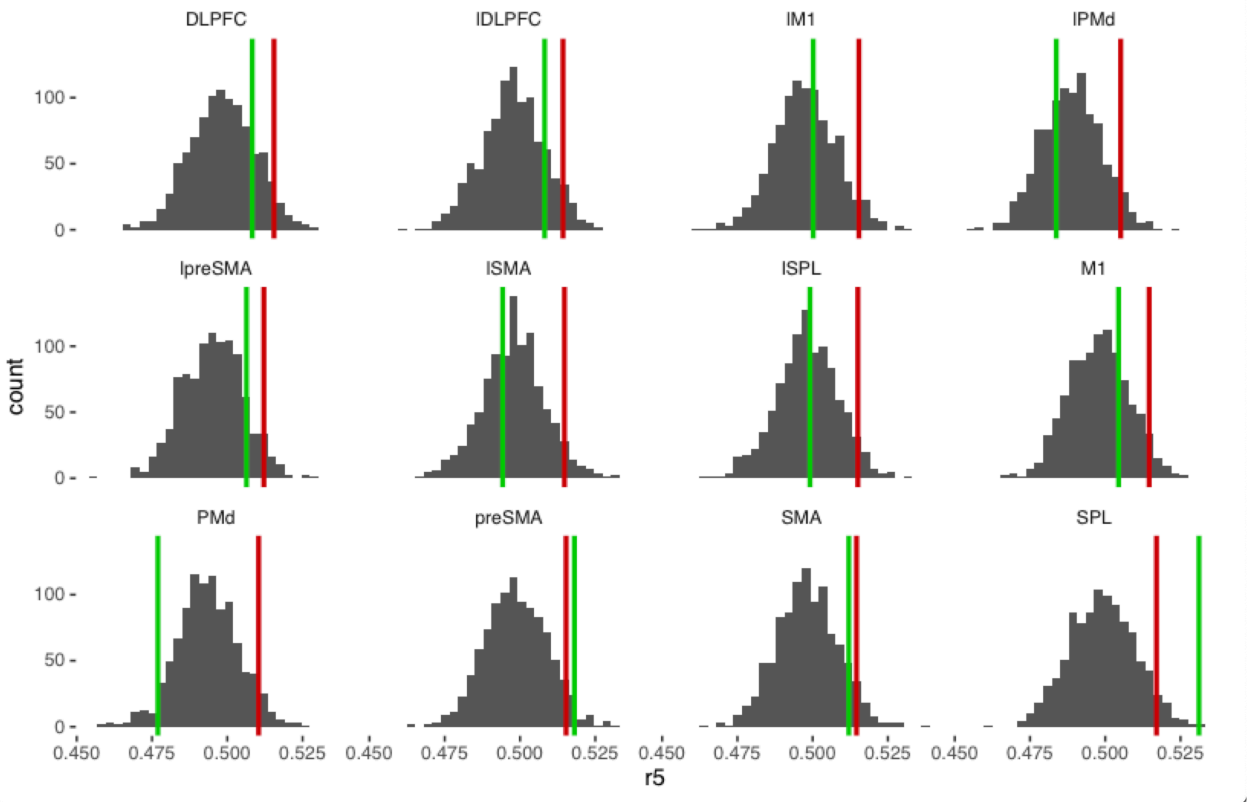

**Figure S2B.** *$10 ROI Decoding vs chance.*

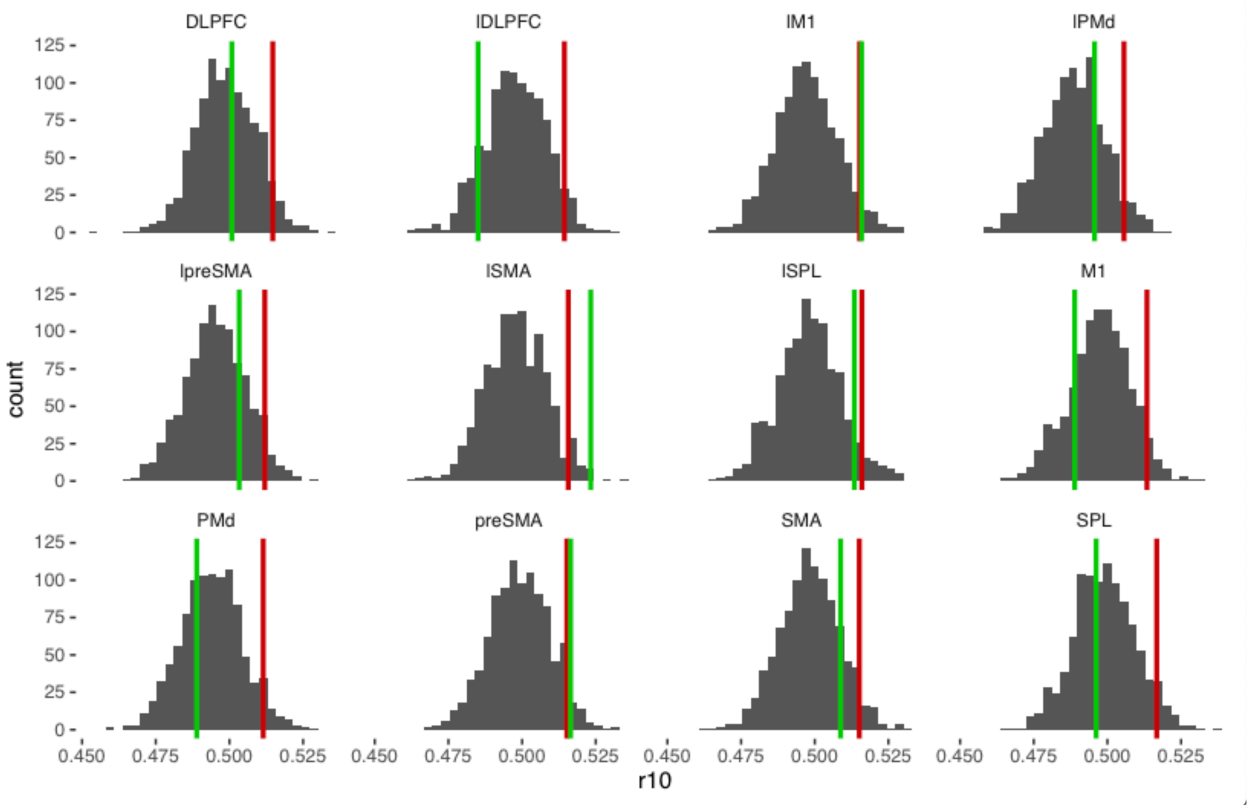

**Figure S2C.** *$30 ROI Decoding vs chance.*

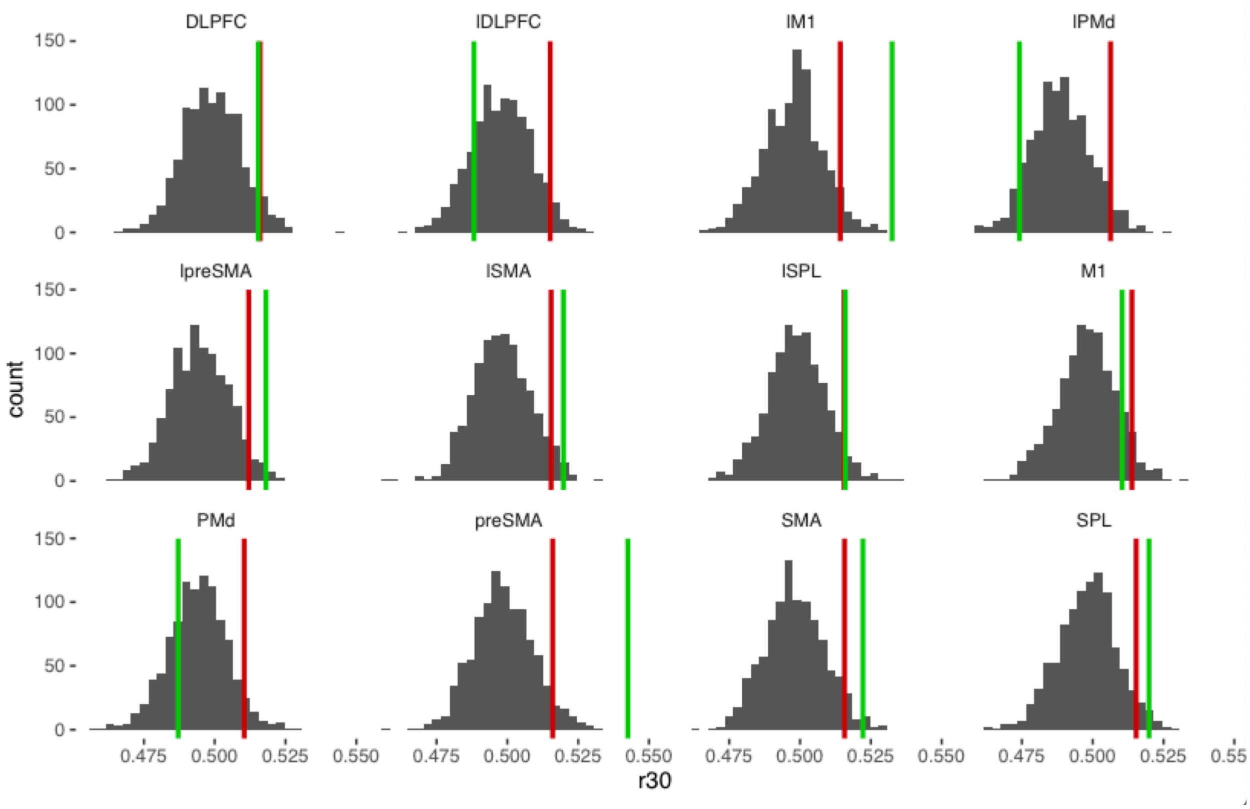

Supplement

**Figure S2D.** *$30 – $5 ROI Decoding vs chance.*

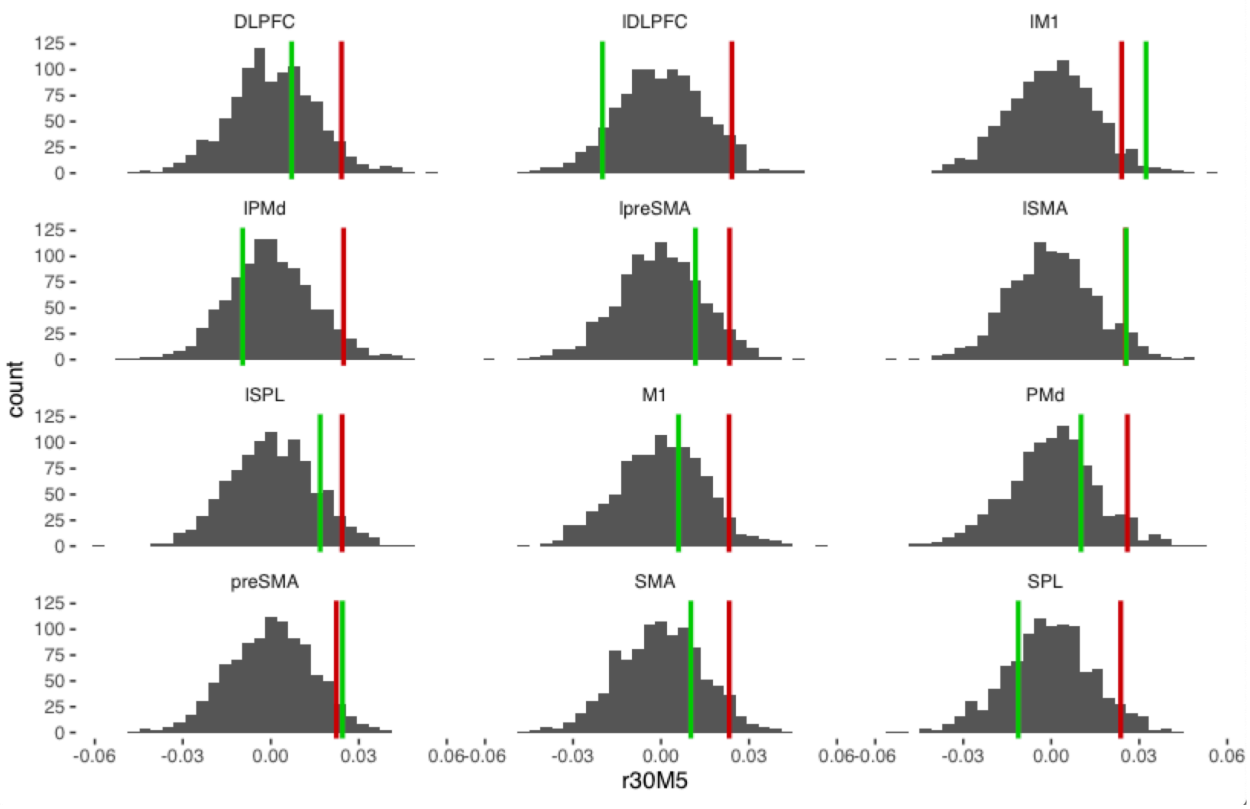

**Figure S2E**. *$30 – $10 ROI Decoding vs chance.*

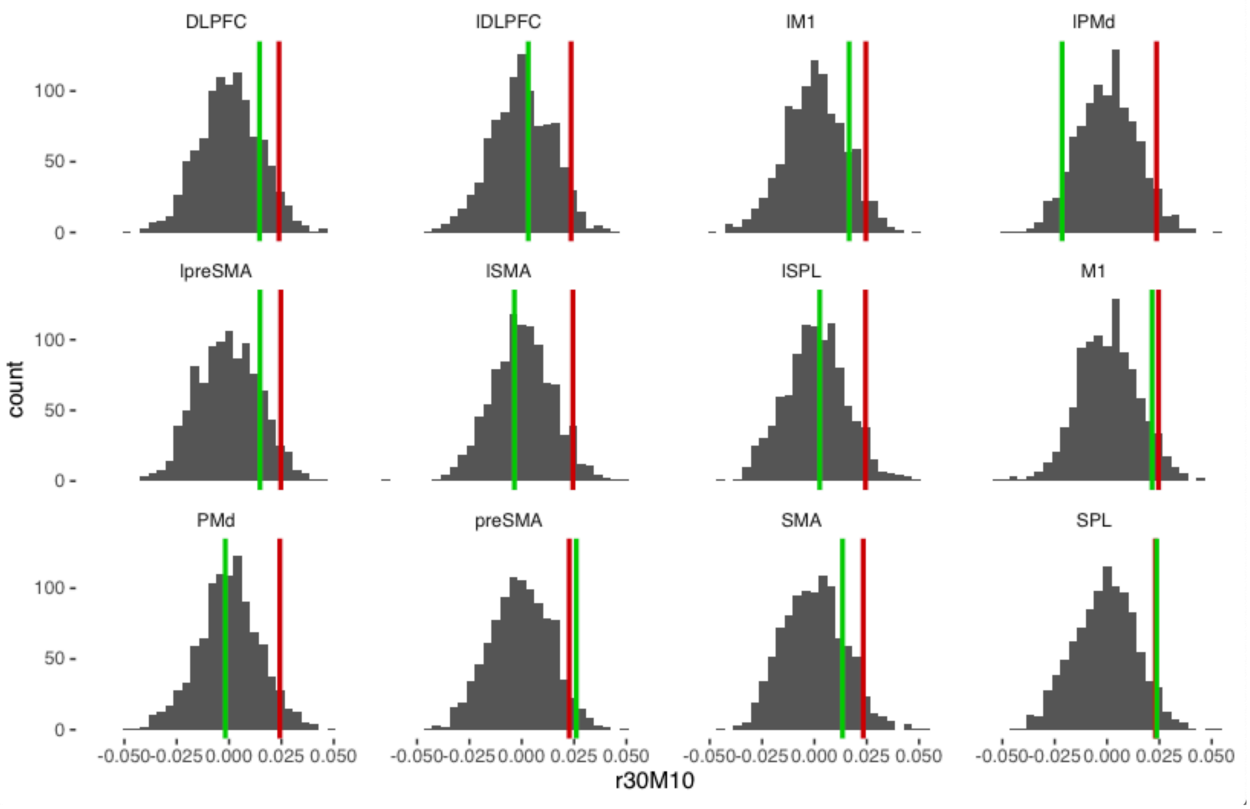

**Figure S3A.** *Effect of reward on action decoding for each subject.* These are the subject-level means used to compute the group-level means shown in Figure 7A from the main text.

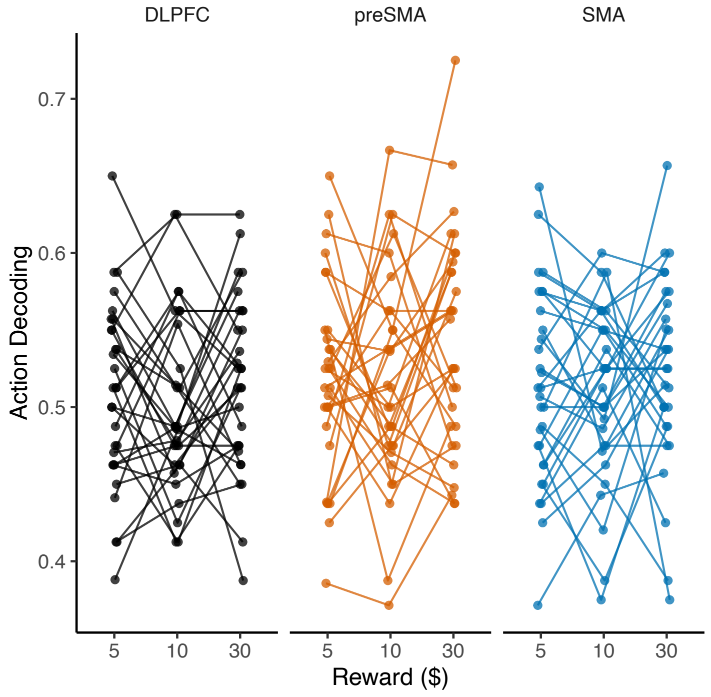


**Figure S3B.** *Effect of action decoding on action performance for each subject.* Thes are the subject-level means used to compute the group-level means shown in Figure 7B from the main text.

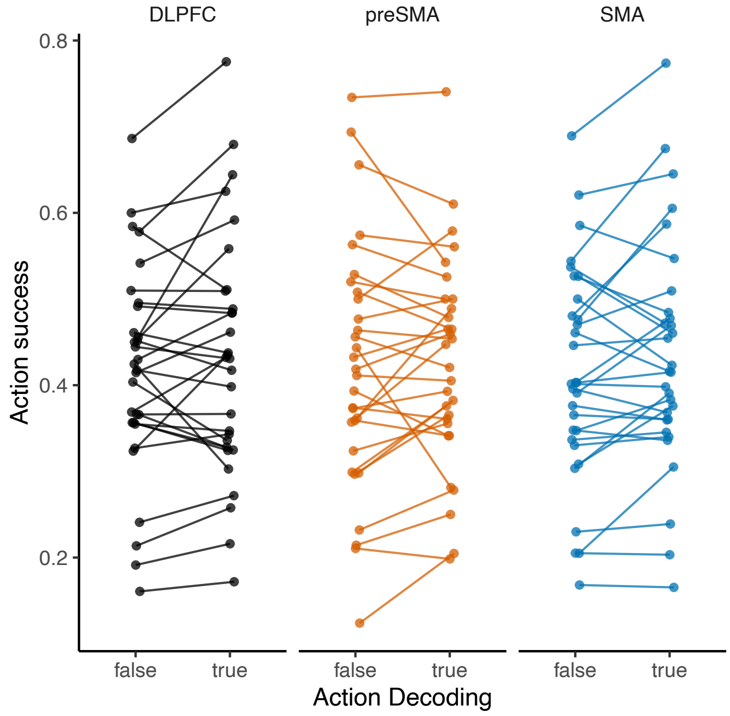

**Figure S4.** *Movement time limits for each subject and sequence and trial of the test session. For participants 104 and 105, an early run was aborted because of miscommunication between the scanner and the stimulus computer. Time limits reverted to their initial value following the resumption of the task.*

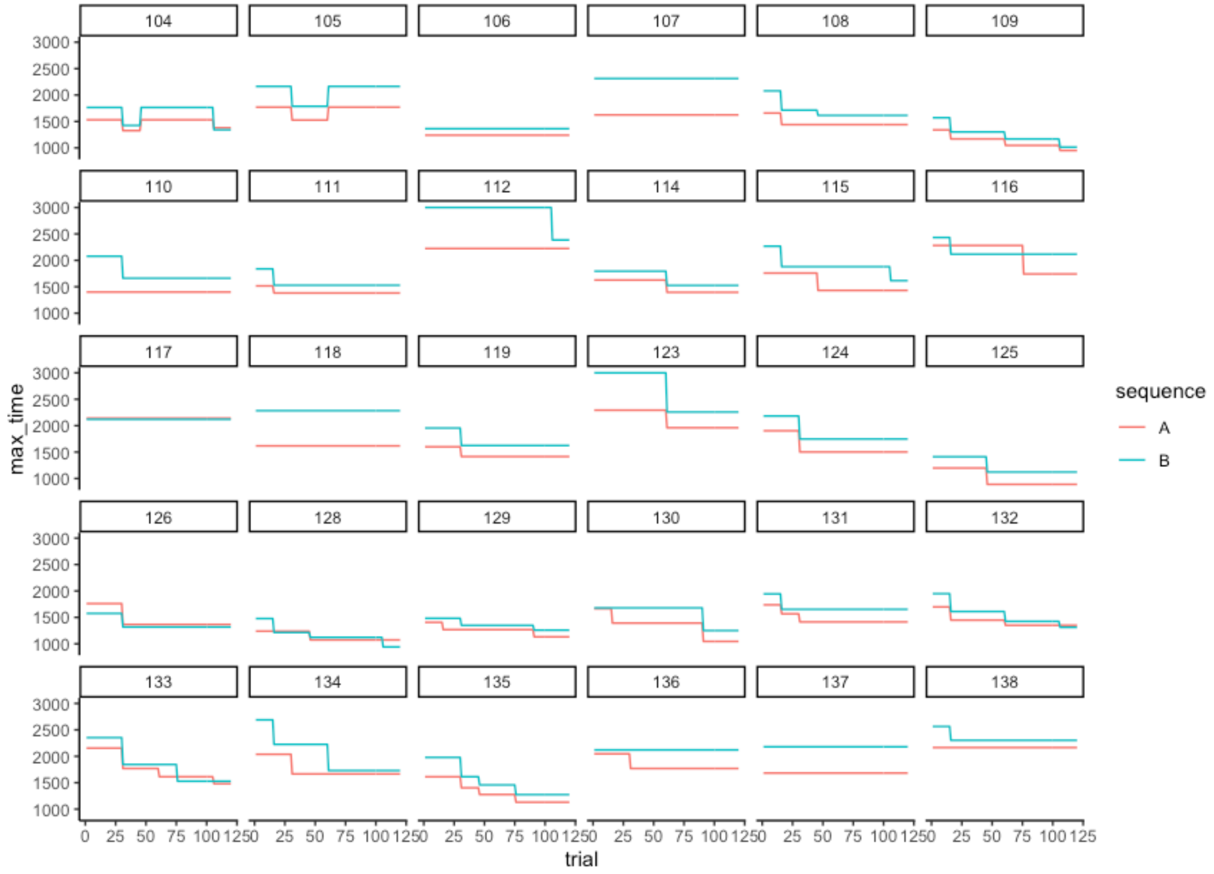


**Figure S5.** *Bootstrapped distributions of group mean SpaceNet classification accuracies.*

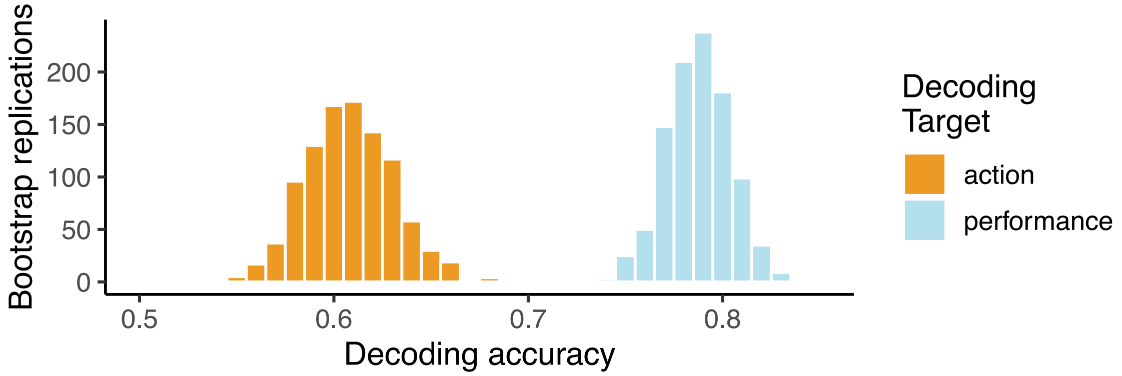

**Figure S6.** *Family-wise error rate analysis for ROI permutation tests.* To examine whether our statistically significant decoding results could have been family-wise errors, we used 1000 bootstrap samples from the null distributions and computed the proportion of bootstrap samples in which one, two, three, four, five, or six ROIs showed a statistically significant effect (p < .05 for their respective distributions). For this analysis, we included six ROIs: DLPFC, preSMA, SMA, PMd, M1, and SPL. The results show that it was unlikely (p < .05) under the null to obtain consistent significant effects across multiple ROIs, as we report in the main text.

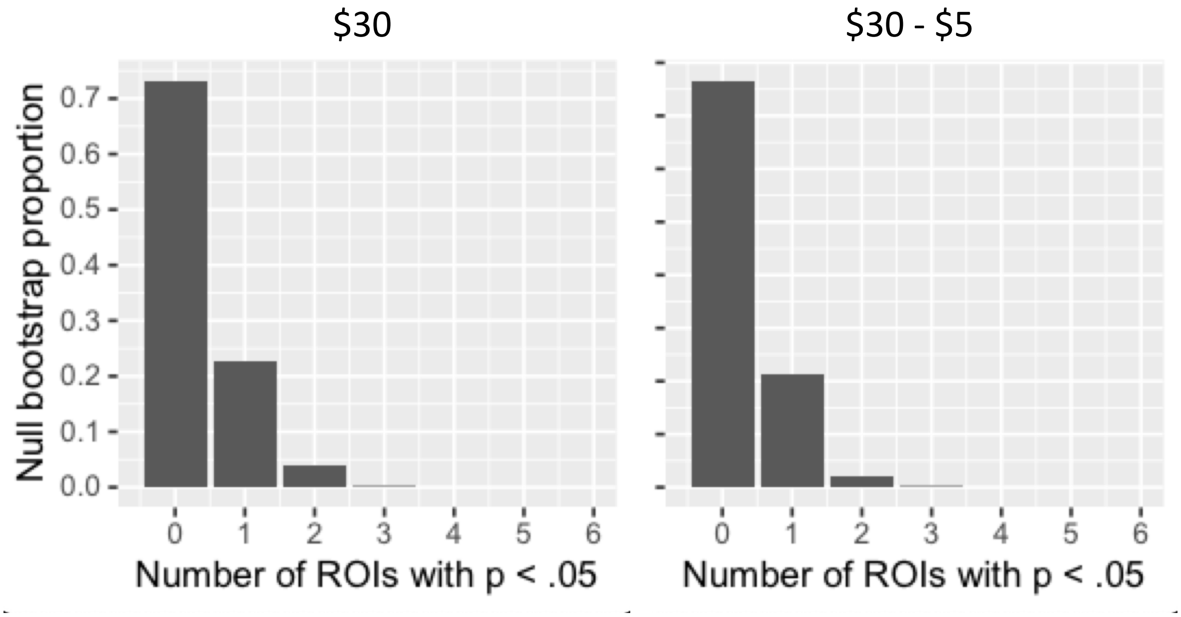


**Figure S7.** *Comparison of reward and action maps.* In the first row are slices showing areas whose activity increased with reward size at cue. In the second row are slices showing the subset of this reward-modulated map that also carried information about action. The maps are similar because the second is a subset of the first, however the action decoding map is much sparser by comparison.

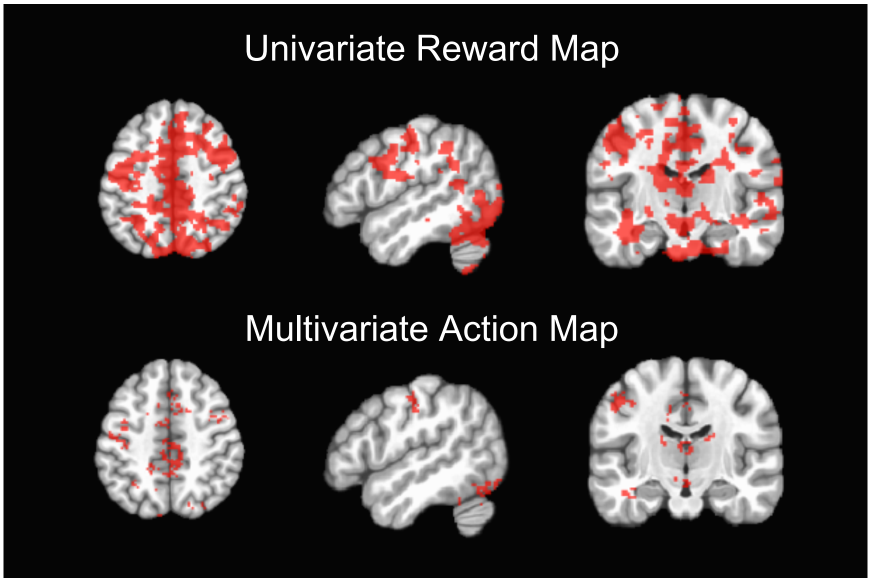

Supplement

**Table S1** *Reward modulation map clusters, corresponding to Figure 2.* The initial group mask contained a single cluster with hundreds of thousands of voxels, so we eroded by 2 voxels to provide some regional clarity and report these clusters below. This table was created using afni's 3dclusterize and wherami commands with an MNI template.

| #Voxels | CM x | CM y | CM z | Peak x | Peak y | Peak z | Overlapping Anatomical ROIs |
|---|---|---|---|---|---|---|---|
| 1351 | −1.9 | +79.9 | +5.3 | −11.0 | +87.0 | −17.0 | B Calcarine Gyrus, R Lingual Gyrus |
| 628 | −6.3 | +59.7 | +48.7 | −15.0 | +77.0 | +31.0 | B Precuneus, R Middle Frontal Gyrus |
| 578 | −3.2 | −16.6 | +41.1 | +1.0 | −21.0 | +27.0 | B Middle Cingulate Cortex, SMA |
| 413 | +34.7 | +73.6 | −19.8 | +21.0 | +89.0 | −27.0 | L Cerrebellum (Crus 1) |
| 199 | −1.3 | +34.8 | +4.8 | +1.0 | +47.0 | −1.0 | Cerebellar Vermis (4/5) |
| 180 | −40.3 | +68.2 | −18.1 | −43.0 | +71.0 | −23.0 | R Fusiform Gyrus |
| 139 | −57.6 | +43.9 | +26.9 | −57.0 | +45.0 | +17.0 | R Supramarginal Gyrus |
| 84 | −47.4 | −15.8 | +24.9 | −45.0 | −13.0 | +21.0 | R Inferior Frontal Gyrus |
| 77 | +41.5 | −4.6 | +30.4 | +43.0 | −7.0 | +25.0 | L Precentral Gyrus |
| 64 | −43.9 | −5.4 | +42.7 | −43.0 | −7.0 | +35.0 | R Precentral Gyrus |

**Table S2** *Sequence information map clusters, corresponding to Figure 3B.* The initial group mask contained a large cluster of 5k+ voxels, so we used a nearest-neighbors setting of 1 to obtain the clusters below.

| #Voxels | CM x | CM y | CM z | Peak x | Peak y | Peak z | Overlapping Anatomical ROIs |
|---|---|---|---|---|---|---|---|
| 1778 | −6.0 | +77.9 | +12.0 | −3.0 | +77.0 | −19.0 | B Calcarine Gyrus, R Lingual Gyrus |
| 1773 | +1.3 | +15.0 | +46.7 | −3.0 | −21.0 | +23.0 | B Middle Cingulate Cortex, Left Precuneus |
| 597 | +39.1 | +72.5 | −17.9 | +45.0 | +63.0 | −31.0 | L Fusiform Gyrus, L Cerebellum (Crus 1) |
| 447 | −2.3 | +34.1 | +3.1 | −11.0 | +31.0 | −9.0 | R Lingual Gyrus, Cerebellar Vermis (4/5) |
| 319 | +6.2 | +7.2 | +13.6 | −7.0 | −3.0 | +3.0 | L Thalamus, L Caudate Nucleus |
| 254 | −58.1 | +42.0 | +25.2 | −59.0 | +43.0 | +15.0 | R SupraMarginal Gyrus |
| 228 | −40.9 | −19.9 | −2.4 | −43.0 | −13.0 | −11.0 | R Insula Lobe |
| 217 | −35.5 | +74.2 | −17.8 | −39.0 | +85.0 | −23.0 | R Fusiform Gyrus |
| 203 | −47.8 | +60.6 | −19.5 | −47.0 | +65.0 | −31.0 | R Inferior Temporal Gyrus |
| 184 | −51.3 | +27.9 | −10.4 | −53.0 | +23.0 | −23.0 | R Middle Temporal Gyrus |

**Table S3** *Performance information map clusters, corresponding to Figure 4B.* Since this map was sparser than the others, we used a more liberal nearest neighbor setting of 3 to obtain the clusters below.

| #Voxels | CM x | CM y | CM z | Peak x | Peak y | Peak z | Overlapping Anatomical ROIs |
|---|---|---|---|---|---|---|---|
| 481 | −4.1 | +79.2 | +4.7 | −27.0 | +73.0 | −21.0 | R Lingual Gyrus, B Calcarine Gyrus |
| 152 | −15.6 | +74.1 | +43.5 | −11.0 | +79.0 | +25.0 | R Cuneus, R Precuneus |
| 147 | −2.2 | +32.5 | +4.4 | −9.0 | +35.0 | −7.0 | Cerrebellar Vermis (4/5) |
| 107 | +0.9 | −8.2 | +42.4 | +1.0 | −19.0 | +31.0 | B Middle Cingulate Cortex, L SMA |
| 101 | −4.1 | +32.5 | +48.6 | −5.0 | +37.0 | +41.0 | B Middle Cingulate Cortex, R Precuneus |
| 58 | −47.6 | −16.4 | +22.9 | −51.0 | −13.0 | +15.0 | R Inferior Frontal Gyrus |
| 53 | +11.0 | +52.6 | +58.7 | +7.0 | +47.0 | +51.0 | L Precuneus |
| 43 | +36.2 | +64.2 | −18.1 | +49.0 | +61.0 | −27.0 | L Fusiform Gyrus, L Cerebellum (VI) |
| 40 | −5.4 | −23.9 | +29.2 | −7.0 | −21.0 | +25.0 | R Anterior Cingulate Cortex |
| 37 | −42.1 | +62.2 | −19.4 | −51.0 | +59.0 | −23.0 | R Fusiform Gyrus, R Inferior Temporal Gyrus |

**Table S4** *Conjunction information map clusters, corresponding to Figure 5.* Since this map was also quite sparse, we again used nearest neighbor setting of 3 to obtain the clusters below.

| #Voxels | CM x | CM y | CM z | Peak x | Peak y | Peak z | Overlapping Anatomical ROIs |
|---|---|---|---|---|---|---|---|
| 157 | −4.6 | +77.1 | +2.8 | −15.0 | +77.0 | −11.0 | B Calcarine Gyrus, B Lingual Gyrus |
| 53 | −4.5 | +30.9 | +1.0 | −7.0 | +33.0 | −5.0 | R Lingual Gyrus, R Thalamus |
| 45 | −18.6 | +73.8 | +43.3 | −23.0 | +73.0 | +35.0 | R Precuneus, R Superior Occipital Gyrus |
| 30 | −2.8 | +35.7 | +48.3 | +5.0 | +35.0 | +43.0 | B Middle Cingulate Cortex |
| 24 | −4.2 | −11.7 | +43.5 | +1.0 | −15.0 | +39.0 | B Middle Cingulate Cortex, B SMA |
| 18 | −5.2 | −24.0 | +29.7 | −3.0 | −19.0 | +27.0 | R Anterior Cingulate Cortex |
| 15 | +4.1 | +42.6 | +1.4 | +3.0 | +43.0 | −3.0 | Cerebellar Vermis (4/5), L Cerebellum (IV−V) |
| 15 | −44.7 | −5.4 | +40.6 | −45.0 | −7.0 | +37.0 | R Middle Frontal Gyrus, R Precentral Gyrus |
| 14 | −10.9 | +89.9 | +9.0 | −15.0 | +89.0 | +3.0 | R Calcarine Gyrus |
| 12 | +35.3 | +63.7 | −20.5 | +33.0 | +63.0 | −23.0 | Left Cerebellum (VI) |
